# The evolution of anisogamy does not always lead to male competition

**DOI:** 10.1101/2020.12.18.423382

**Authors:** Mattias Siljestam, Ivain Martinossi-Allibert

## Abstract

Anisogamy has evolved in a large proportion of sexually reproducing multicellular organisms allowing the definition of the female and male sexes, producing large and small gametes, respectively. Anisogamy is the initial sexual dimorphism: it has lead the sexes to experience selection differently, which makes it a good starting point to understand the evolution of further sexual dimorphisms. For instance, it is generally accepted that anisogamy sets the stage for more intense intrasexual competition in the male sex than in the female sex. However, we argue that this idea may rely on assumptions on the conditions under which anisogamy has evolved in the first place. We consider here two widely accepted scenarios for the evolution of anisogamy: *gamete competition* or *gamete limitation*. We present a mechanistic mathematical model in which both gamete size and an intrasexual competition trait for fertilisation can coevolve in a population starting without dimorphism between its two mating types. Two different intrasexual competition traits are investigated, gamete motility and the ability of gametes to capture gametes of the opposite mating type. We show that *gamete competition* and *gamete limitation* can lead to greatly different outcomes in terms of which sex competes most for fertilisation. Our results suggest that *gamete competition* is most likely to lead to stronger competition in males. On the other hand, under *gamete limitation*, competition in form of motility can evolve in either sex while gamete capture mainly evolves in females. This study suggests that anisogamy does not *per se* lead to more intense male competition. The conditions under which anisogamy evolves matter, as well as the competition trait considered.

---

Most sexually reproducing multicellular organisms have evolved mating types and anisogamy (dimorphism in gamete size)(Lessells et al., 2009). In systems with two mating types, the one producing larger gametes is denoted the female sex while the one producing small gametes is the male sex. Because it is the earliest possible sexual dimorphism, researchers interested in sexual selection have tried to understand how anisogamy may have influenced the evolution of sexual dimorphisms or sex-biases that evolved later on, most notably in intrasexual competition for mating and in parental care. Two main approaches have been used to this end: first the study of sex-biases in intrasexual competition and parental care in contemporary species through empirical observation, supported by meta-analysis and review (Clutton-Brock, 1991; Cox and Calsbeek, 2009; Janicke et al., 2016; Singh and Punzalan, 2018) and second mathematical modelling (e.g. Kokko and Rankin, 2006; Kokko et al., 2012; Jennions and Fromhage, 2017). The empirical approach is most powerful to inform us on how ecological settings, evolutionary history, and particularities of the mating system may influence sexual dimorphisms in competition and parental care, but it does not prove very insightful when it comes to revealing the influence of anisogamy on these traits because of the multitude of confounding factors that are not entirely understood. On the other hand, the theoretical approach on which we will focus relies on thought experiments and mathematical models that aim at understanding the evolutionary origins of the male and female sexes, i.e. the evolution of anisogamy. This line of research has also endeavoured to generate predictions on further evolutionary changes that would result from the state of anisogamy, sometimes referred to as the sexual cascade (Parker, 2014), where anisogamy ultimately leads to sexual dimorphisms in intrasexual competition and may also influence dimorphism in parental care. We believe this approach to be most useful to specifically understand the influence of anisogamy on further sexual dimorphisms, because it starts in a context where anisogamy is the only difference between males and females, therefore automatically removing the influence of potentially confounding factors.

Starting in 1948, Bateman (Bateman, 1948) conducted an influential and later questioned and debated (Sutherland, 1985; Tang-Martinez and Ryder, 2005; Gowaty et al., 2012) study in *Drosophila melanogaster* in which he measured higher variance in reproductive success in males than in females. He suggested that this result may be explained by the state of anisogamy in the following way: males produce small, cheap, numerous gametes and females fewer, larger, more energetically costly ones, therefore the number of gametes produced by males would never be a limiting factor of their reproductive success, while it is the case for females. This would allow males to reach higher potential reproductive success than females. As a consequence, because each offspring has a male and female parent and the population sex-ratio is assumed to be balanced, the fact that some males reach a high reproductive success implies that other males will have low or null reproductive success. At the same time, most females are expected to have a reproductive success that is close to average. This results in a higher variance in reproductive success for males. In the early 1970s, Trivers (Trivers et al., 1972) followed up on Bateman’s idea and suggested an explanation for the causal relationship between anisogamy, intrasexual competition and parental investment. Trivers argued that initial energy investment in offspring should determine which sex competes and which sex cares for offspring. The larger initial energy investment in the zygote by females should automatically trigger competition for mating opportunities in males and create a stronger incentive in females to provide further care in order to increase offspring survival. Trivers’s ideas (Trivers et al., 1972) regarding parental investment have been critically examined early on and resulted in the production of a wealth of theoretical work (e.g. Dawkins and Carlisle, 1976; Queller, 1997; Kokko and Jennions, 2008; Jennions and Fromhage, 2017), which have shown that although there may be a causal link between anisogamy and parental investment, it ought to be more complex than Trivers had initially predicted (Knight, 2002). In contrast, the mechanism through which anisogamy should result in more intense intrasexual competition in males has been relatively neglected by theoreticians. Although initially a verbal model, it has not received much critical attention until recently (see Lehtonen et al., 2016), and instead has been supported by verbal arguments (e.g. Schärer et al., 2012; Parker, 2014) or correlational empirical data (Janicke et al., 2016).

Here, we re-examine the causal relationship between anisogamy and the evolution of intrasexual competition. Although much work has been done on the evolution of anisogamy since Trivers developed his ideas (to cite a few: Parker et al., 1972; Parker, 1978; Schuster and Sigmund, 1982; Smith, 1982; Hoekstra et al., 1984; Hoekstra, 1984; Bulmer, 1994; Dusenbery, 2000; Bulmer and Parker, 2002; Dusenbery, 2006; Lessells et al., 2009; Lehtonen and Kokko, 2011), the interaction between the evolution of gamete size and the evolution of intersexual competition traits is still not well understood. We argue that considering the coevolution of anisogamy and intrasexual competition (as opposed to their independent or sequential evolution) is important, because the selection pressures that are responsible for the evolution of anisogamy may still be at play when intrasexual competition evolves. For this reason we develop a model where both gamete size and a competition trait are evolving simultaneously, starting from a state of isogamy (two mating types with equal gamete size). Classically, there are two main theoretical frameworks for the evolution of anisogamy (Lehtonen and Parker, 2014), *gamete competition* and *gamete limitation* (detailed below) and we want to examine whether these should favour different sexual dimorphisms in intrasexual competition.

Most models of the evolution of anisogamy assume that there is a limited amount of energy an individual may allocate to gamete production resulting in a trade-off between gamete number and individual gamete size (for example: Parker et al., 1972; Smith, 1982; Bulmer and Parker, 2002). In the *gamete competition* framework, initially developed by Parker (Parker et al., 1972), the evolution of anisogamy is mainly driven by a constraint imposed on the survival of the zygote (Bulmer and Parker, 2002), which depends on gamete size. Larger gametes harbour more resources and provide zygotes with higher survival chances. Once one mating type starts to evolve larger gametes, the other mating type is somewhat released from the constraint of providing energy to increase zygote survival and can therefore reduce energy investment into individual gametes in order to increase the number of gametes produced, which is a good strategy in this context to increase fertilisation success. In this framework, there is no sperm limitation because gamete density is high enough and gametes encounter each other easily, which means that all larger gametes are automatically fertilised while some proportion of the smaller gametes are in excess. This situation is thus reminiscent of the view that Bateman and Trivers held, which is that female reproductive success is limited by female fecundity, while male reproductive success is limited by male competitive ability. We therefore predict, in agreement with previous work (Schärer et al., 2012; Lehtonen et al., 2016), that if anisogamy evolves in a *gamete competition* context, it should favour the evolution of competition traits in the male sex.

In the *gamete limitation* framework, which originated with Kalmus (Kalmus, 1932) the evolution of anisogamy arises as a strategy to increase gamete encounter rates, in an environment where gamete density is very low. The evolution of anisogamy in this framework is favoured by sperm limitation, i.e. the fact that not all the larger gametes get fertilised and this suggests that both sexes experience selection to increase fertilisation success. For that reason, we predict that the *gamete limitation* context should allow the evolution of competition traits in both sexes.

As demonstrated by Lehtonen and Kokko (Lehtonen and Kokko, 2011), *gamete competition* and *gamete limitation* are not two mutually exclusive theories on the evolution of anisogamy, but rather two complementary ideas that speculate on how anisogamy may have evolved in populations of high or low gamete densities. It is therefore relevant to include both scenarios in the same model, which we do here by including gamete density as a variable parameter of our model (regulated by population density or the per individual gamete energy allocation, or both). Furthermore, *gamete limitation* may be a particularly relevant scenario since the organisms that are thought to have evolved anisogamy in the first place are sessile broadcast spawning marine invertebrate (Lehtonen and Parker, 2014; Parker, 2014), and this type of organism can easily be subjected to sperm limitation (Levitan and Petersen, 1995; Levitan, 1998a,b; Yund, 2000; Crean and Marshall, 2008).

In the present study, we develop and analyse a mathematical model of the coevolution of gamete size together with an intrasexual competition trait, starting from a population without sexual dimorphism. By varying gamete density, we explore several scenarios ranging from extreme *gamete limitation* (low density) to intense *gamete competition* (high density). Intrasexual competition traits in this model are gamete-level traits that increase the fertilisation success of an individual at the expense of the success of other individuals of the same mating type. We investigate independently the evolution of two different competition traits. The first one is gamete motility, which in nature is often found prominently in the smaller gametes. The second trait is the ability of gametes to capture nearby gametes of the opposite mating type in order to achieve fertilisation by increasing their apparent size, a strategy observed in nature under the form of a jelly coat surrounding the larger gametes of several marine invertebrates (e.g. Farley and Levitan, 2001; Podolsky, 2002). We explore a wide range of possible scenarios and our model provides novel insights on the relationship between the evolution of anisogamy and sex-biases in intrasexual competition.

## Model

Here, we present a mechanistic mathematical model of the evolution of anisogamy, where we consider the coevolution of gamete size and an intrasexual gamete competition trait for fertilisation. We consider a community of sessile marine animals reproducing through external fertilisation. This is inspired by the conditions under which anisogamy is thought to have first evolved in animals (Parker, 2014). Below, we first give a verbal presentation of the structure and analysis of the model before going into the mathematical details.

The life cycle of the organism is modelled in two steps and assumes discrete non-overlapping generations. In the first step of the life cycle gametes are produced. A fixed number of brooding spots on the sea floor are occupied by individuals (zygotes) that produce gametes. The number of gametes produced per zygote is traded off against both gamete size and the proportion of energy spent on an intrasexual gamete competition trait, with larger gametes having a higher chance of surviving to next step of the life cycle (Figure 1). In the second step of the life cycle gametes fertilisation occurs. The surviving gametes are synchronously released into a common mating pool, and fertilisation is the result of collisions of gametes of the two mating types, generating the zygotes of the next generation (Figure 2). These newly formed zygotes have to survive to enter the next generation with larger zygotes having a higher chance of survival (Figure 1).

**Fig. 1.**
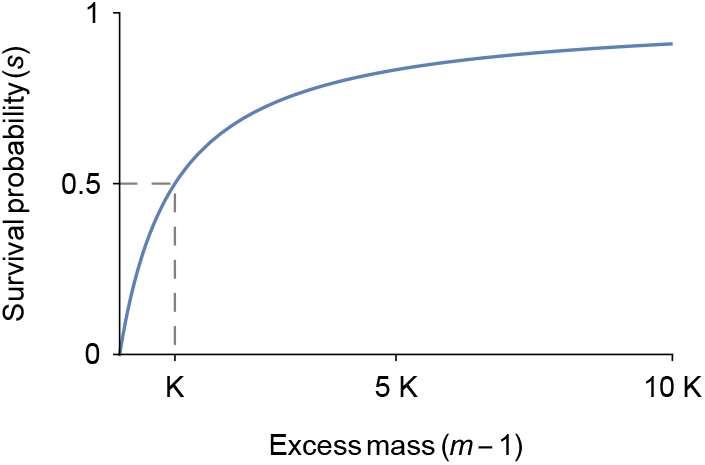
Survival probability of gametes and juvenile zygotes as a function of their excess mass allocated to survival, *m* – 1. A survival of 50% is reached when the excess mass equals the half-saturation constant *K* marked out by the dashed line.

**Fig. 2.**
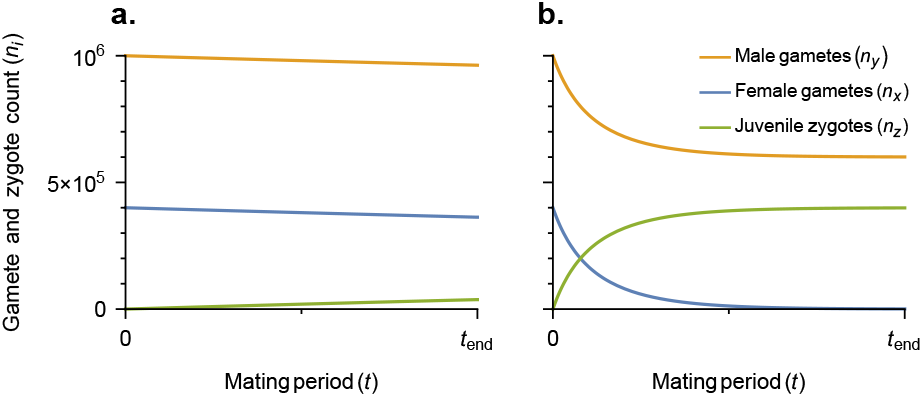
Number of gametes and newly formed juvenile zygotes per adult zygote over the mating period under **a.** *gamete limitation* (low gamete density) and **b.** *gamete competition* (high gamete density). These graphs are given by equation A6 with parameters: initial female gamete count *n*_*x*, 0_ = 5 × 10^5^, initial male gamete count *n*_*y*, 0_ = 10^6^ and the product of *pdσ_xy_ v_xy_* (which is proportional to the gamete fertilisation rate) equals 10^−5^ in **a.** and 10^−7^ in **b.**

In our analysis, we investigate the evolution of anisogamy by allowing for coevolution of two gametic traits: gamete size and the relative investment into a competition trait. We assume evolution to start under the constraint of isogamy, meaning equal trait values for both mating types. This constraint can be removed by disruptive selection. We first solve for isogamic attractors under constrained isogamy (i.e. gamete trait values to which evolution leads). Then, we determine if these isogamic attractors are endpoints of evolution or if disruptive selection appears leading to the evolution of anisogamy. When anisogamy arises, we then use simulations to determine how the trait values of gamete size and investment in competition for the two mating types evolve. By varying gamete density in the model we transition from analysing the evolution of anisogamy in a *gamete limitation* context at low density to a *gamete competition* context at high density.

### Gamete production

In the first step of the life cycle, zygote individuals are randomly chosen to occupy brooding spots occurring with a density of 2*d*, where the zygotes then can grow and produce gametes. The zygotes are assumed to be in abundance such that all brooding spots are occupied. The two mating types *x* and *y* are of equal frequency such that zygotes of each mating type occupies spots with a density of *d*.

Each zygote possesses a fixed amount of energy *e* allocated to gamete production. A proportion of the energy *r_i_* is spent on a gamete competition trait (see details further down), where *i* denotes the mating type and can be either *x* or *y*. We refer to the trait *r_i_* as relative competition investment. The remaining energy *e*(1 – *r_i_*) is converted into gamete mass with gametes of size *m_i_*. The number of gametes *n_i_* produced per zygote is thereby a function of both the gamete size *m_i_* and the relative competition investment *r_i_* and is given by

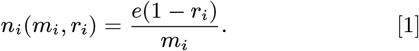

The probability that a gamete survives until it enters the mating pool (the second step of the life cycle) is an increasing function of its size *m_i_*. We assume gamete cells must have a minimum size of 1 to perform their function and that any excess mass, *m_i_* – 1, represents resources provisioned for survival. The survival probability of a gamete *s_g_* is given by following saturating function *s_g_*(*m_i_*) = (*m_i_* – 1)/(*K_g_* + *m_i_* – 1), where *K_g_* gives the half-saturation constant of gamete survival, i.e. the excess mass required for a 50% probability of survival (Figure 1). We will refer to *K_g_* as the gamete survival constraint.

Hence, the number of gametes per zygote of mating type *i* entering the mating pool, *n*_*i*, 0_, equals the number of gametes produces times their survival probability: *n*_*i*, 0_(*m_i_*, *r_i_*) = *n_i_*(*m_i_*, *r_i_*)*s_g_* (*m_i_*).

### Gamete fertilisation

In the mating pool, the second and last step of the life cycle, gamete fertilisation occurs, which produces the zygotes of the next generation. Here, the gametes that survived enter the mating pool where they move around randomly and can collide with each other. Each collision between gametes of the two mating types *x* and *y* can result in a fertilisation with probability *p*. A fertilisation event removes both gametes from the mating pool and produces a juvenile zygote offspring. To determine the collision frequency between gametes of the two mating types we use collision theory (originally used to model chemical reaction rates between gas particles, McNaught et al., 2014). The collision frequency depends on three factors: the sizes of the gametes *m_x_* and *m_y_*, the gamete speeds *v_x_* and *v_y_* (see details further down) and the density of gametes, where all three factors increases the collision frequency (See in Appendix 1, Eq. A1).

The initial gamete density of mating type *i* is given by *n*_*i*, 0_(*m_i_*, *r_i_*)*d*, i.e. the number of surviving gametes per zygote entering the mating pool (at *t* = 0) multiplied by the zygote density *d*. The number of gametes then declines over time as they collide and get fertilised (Figure 2, Eq. A6).

Finally, the probability that a newly formed juvenile zygote survives until the next generation is an increasing function of its size *m_z_* (which is given by the combined sizes of the two gametes that fused to produce it *m_z_* = *m_x_* + *m_y_*) according to *s_z_* (*m_x_*, *m_y_*) = (*m_x_* + *m_y_* – 1)/(*K_z_* + *m_x_* + *m_y_* – 1), where *K_z_* gives the half-saturation constant of zygote survival (Figure 1), and we will refer to *K_z_* as the zygote survival constraint.

### The intrasexual competition traits

We investigate separately two alternative intrasexual competition traits: gamete motility or fusion partner capture.

#### (i) Competition through motility

In this scenario we assume that gametes are in still water resulting in no basal motility of the gametes. The competition trait is then a flagella-like mobility structure that allows gametes to move and encounter a fusion partner.

The amount of energy spent on the competition trait per gamete is given by *m_i_r_i_*/(1 – *r_i_*) (see Eq. 1). To determine the gamete speed *v_i_* we add three parameters to this expression:

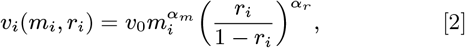

where *α_m_* and *α_r_* determine how gamete speed is affected by gamete size *m_i_* and the relative investment *r_i_*, respectively (with *α_r_* > 0). *v*_0_ is a velocity scale factor.

We refer to *α_r_* as competition investment efficiency. It describes how profitable it is to invest more energy into the competition trait rather than gamete mass. Whenever *α_r_* < 1, there is diminishing rate of return in speed *v_i_* with increasing *r_i_*, while *α_r_* = 1 results in a linear return in speed and *α_r_* > 1 results in increasing rate of return with increasing investment *r_i_*. We refer to *α_m_* as the size-dependent (competition) investment efficiency, which determines whether there is a positive or negative relationship between gamete size and return from investing into the competition trait. When *α_m_* > 0, larger gametes are faster than smaller gametes for a given proportional investment *r_i_*, while *α_m_* < 0 results in smaller gametes being faster for the same *r_i_*. These two investment efficiency parameters give flexibility in the description of the competition trait, allowing the model to cover a variety of possible scenarios.

#### (ii) Competition through fusion partner capture

In this scenario, we assume that all gametes are in suspension in water currents, and have similar motility due to the movement of their environment with gamete speed given by just the basal velocity *v*_0_. The competition trait is here represented by an extension of apparent gamete size (for example by producing a jelly coat around the gamete, or filaments extending from the gamete) which allows to capture potential fusion partners (see e.g. Tilney and Jaffe, 1980). This trait is modelled as increasing the apparent gamete size 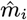. This means that gamete size is effectively increased in terms of collision target size (Eq. A1), but not in terms of survival. Apparent size is defined as

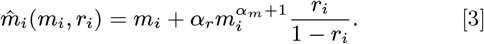

The second term gives the increase in apparent size due to investment in competition, and can be divided into two products. First *m_i_*(*r_i_*/(1 – *r_i_*), giving energy invested into the increased apparent size and second 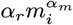, giving how many times cheaper the artificial mass is compared to increasing the actual gamete size *m_i_*. For this trait we also refer to *α_r_* and *α_m_* as competition investment efficiency and size-dependent investment efficiency, respectively.

## Analysis

We investigate the coevolution of two traits, gamete size *m_i_* and relative competition investment *r_i_*, in the two mating types *x* and *y*. We assume these traits to be coded by autosomal chromosomes, such that the traits of a genotype is characterised by the vector *T* = (*m_x_, r_x_, m_y_, r_y_*) giving the two traits of the two mating types, to which we refer as the strategy of the genotype. Note that each individual will only express the two traits corresponding to its own mating type. Initially, we assume evolution to be under isogamic constraint, meaning that the two mating types *x* and *y* have equal gamete size *m_x_* = *m_y_* = *m_c_* and equal relative competition investment *r_x_* = *r_y_* = *r_c_*. Under this isogamic constraint, the strategy of a genotype is thereby given by only these two traits *T*_c_ = (*m_c_, r_c_*), where c denotes the traits being under the constraint.

In our analysis we consider a large resident population starting with a single strategy *T*, to which we iteratively introduce an initially rare mutant strategy 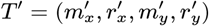 that has a slight deviation in one or more of the traits of the resident strategy *T* (for evolution under isogamic constraint, replace *T* with *T*_c_ and *T*′ with 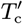). The expected long term growth of a rare mutant strategy *T*′ is given by the fertilisation success of the rare mutant strategy 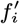 (Eq. A14) divided by the fertilisation success of the resident strategy *f_i_* (Eq. A9), averaged for the two mating types *x* and *y* (Shaw and Mohler, 1953),

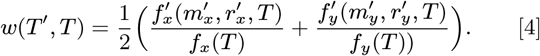

Note that the resident fertilisation success *f_i_* is not affected by the mutant strategy *T*′ as the mutant is assumed to be rare. For the same reason, the mutant fertilisation success 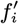 is not affected by the mutant traits of the other mating type. We refer to *w*(*T*′, *T*) as the invasion fitness of the mutant strategy and the mutant is able to increase in frequency and thereby invade whenever *w*(*T*′, *T*) > 1.

### Adaptive dynamics

By assuming rare mutations of small effect we can use the adaptive dynamics framework to predict the evolutionary dynamics (Metz et al., 1992; Dieckmann and Law, 1996; Geritz et al., 1998). Small mutational effect ensures that a mutant strategy that invades will always replace and become the new resident strategy as long as the population is under directional selection (Dercole and Rinaldi, 2008, Appendix B) and rare mutations ensures that a mutant strategy will either fixate or go extinct before a new mutant is introduced. If a mutant strategy fixates it will become the new resident strategy *T* of the population. This results in a trait substitution sequence where the strategy of the population *T* evolves in a step-wise manner for each invading mutant strategy.

The selection gradient *β* gives the direction in the trait space of *T* in which mutants have highest invasion fitness, thus giving the expected direction of evolution. The selection gradient *β* is given by the gradient of the invasion fitness (Eq. 4) when the mutant trait *T*′ is similar to the resident traits *T*

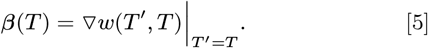

Within the limit of small mutational effect, iterating this mutation-invasion process results in an gradual evolutionary path given by

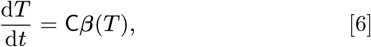

(Dieckmann and Law, 1996; Champagnat et al., 2006; Durinx et al., 2008), where C is the product of the mutational variance-covariance matrix, the mutation rate and population size.

### Predicting the evolutionary outcomes

We can predict the outcomes of the evolutionary path (as given by Eq. 6) using the adaptive dynamics framework. Strategies *T* where directional selection ceases, i.e *β*(*T*) = 0, are of special importance. These strategies are called singular strategies and we denote them *T*\* (or 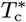 for evolution under the isogamic constraint). A singular strategy *T*\* has two important stability properties. First, *T*\* can be either an attractor or a repeller of the evolutionary dynamics. Determining this tells us whether evolution will approach it or not, and we refer to this property as the *attractiveness*. Second, *T*\* can be invadable or uninvadable by nearby mutant strategies and we refer to this property as the *invadability* (more details are given in Appendix 5.2)(Leimar, 2009).

Each strategy of the isogamic constrained evolution *T*_c_ = (*m_c_*, *r*_c_) has a corresponding isogamic strategy for the unconstrained evolution, namely *T* = (*m_c_, r_c_, m_c_, r*_c_) and we denote this strategy *T*_=_. Also, each singular strategy of the constrained evolution 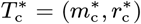 happens to also always be a singular strategy at its corresponding point in the uncon-strained evolution 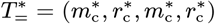 (Van Dooren et al., 2004). This gives each isogamic singular strategy pair 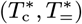 a total of four stability properties: the *attractiveness* and *invadability* of both the constrained and the corresponding unconstrained evolution.

We first allow evolution to occur under the isogamic constraint until it reaches an attractor (singular point) 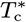. Then, we release the constraint starting evolution from 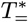 and observe how gamete size and the competition trait in both mating types coevolve. Under this setting we can, as described below, classify the evolutionary dynamics at each singular strategy pair 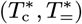 into four different scenarios: it can either be a point that repels isogamic constrained evolution, or it can be an attractor of the constrained evolution in which case three outcomes are possible: evolution transitions from isogamy to anisogamy, evolution comes to a halt, or genetic polymorphism evolves.

#### Classification of the evolutionary dynamics at isogamic singular point

We assume evolution to start under the isogamic constraint and in order to classify the evolutionary dynamics around an isogamic singular strategy pair 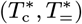 we first look at the *attractiveness* of 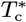, which tells whether the constrained evolution will approach 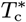 in the first place or not. If 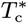 is a repeller it will not be approached by evolution and the other three stability properties (the *invadability* of 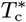 and the *attractiveness* and *invadability* of 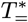) have no relevance.

On the other hand, if 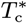 is an attractor, the isogamic constrained evolution will approach it. After 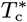 is reached by evolution, two or potentially three different evolutionary outcomes can follow depending on the other three stability properties:

1. An attractor of the isogamic constrained evolution 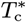 is an evolutionary end-point if both 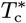 and its corresponding 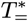 are attracting and uninvadable. Then when evolution reaches 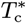 no further mutants can invade in either the constrained or unconstrained trait space and there is no selection for removing the isogamic constraint. Hence, evolution stops and the population stays isogamic.
2. An isogamic attractor 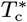 is a point where anisogamy evolves if 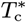 is uninvadable but 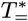 is an invadable repeller for the unconstrained evolution. Then it is a so called saddle point being a fitness maximum in the isogamic constrained trait manifold but a fitness minimum in orthogonal directions (where the gamete traits of the mating types diverge) in the unconstrained trait manifold. This means that as long as the evolution is under the isogamic constraint 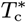 is an attractor that can not be invaded by any further mutants when reached. However, as evolution gets close to 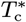 there in an increasing disruptive selection for gamete size *m_i_* and relative competition investment *r_i_* to diverge between the two mating types *x* and *y* for the unconstrained evolution. This disruptive selection acts as a selection pressure for the removal of the isogamic constraint. As soon the the isogamic constraint is removed gamete size *m_i_* and relative competition investment *r_i_* of the two mating types *x* and *y* diverge and the population evolves anisogamy as it repels away from the isogamic strategy 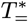. The mating type with larger gametes (arbitrarily) gets the index *x*, and is defined as female while the other mating type *y* is defined as male (*m_x_* > *m_y_*).
3. If an attractor of the isogamic constrained evolution 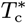 is invadable by nearby mutants it follows that its corresponding unconstrained 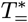 is an invadable repeller (Van Dooren et al., 2004). Here, evolution can lead to two different outcomes. First, because 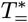 is a repeller of the unconstrained evolution, anisogamy can evolve removing the isogamic constraint, just as described above in case (2). Second, because 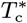 is an invadable attractor for the constrained evolution, it is a special point where isogamic genetic polymorphism can evolve: if 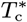 is reached by evolution and the isogamic constraint remains, nearby isogamic mutant strategies can invade and coexist with 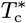 resulting in two isogamic genotype strategies. The strategies of these two isogamic genotypes then diverge as evolution proceeds (evolutionary branching, Geritz et al., 1998). One genotype then evolves larger gametes (with equal size for both mating types) and the other evolves smaller gametes (still equal size for both mating types). Hence, this introduces genetic variation in gamete size in the population becoming pseudo-isogamic, as this gamete size polymorphism within mating types can be seen as an anisogamous situation, but with each genotype remaining isogamous. In summary, at such an invadable isogamic attractor there are two potential evolutionary outcomes: anisogamy if the isogamic constraint is removed, or a pseudo-isogamic polymorphism if the constraint remains.

In this case if the isogamic constraint is removed because anisogamy is favored by selection, then the possibility for isogamic genetic polymorphism is ruled out. There is indeed evidence supporting that the evolution of isogamic polymorphism is unlikely if anisogamy is allowed to evolve, as one might expect anisogamy to evolve first prohibiting the evolution of isogamic polymorphism (Van Dooren et al., 2004). In addition, isogamic polymorphism is a sub-optimal strategy compared to anisogamy in terms of fertilisation success: the most likely type of gamete collision, between a large and a small gamete, has a 50% probability of occurring between gametes of the same mating type and can thus not result in fertilisation. Anisogamy is thereby up to two times more efficient in terms of fertilisation rate. It is therefore possible that if a pseudo-isogamic genetic polymorphism evolves first, it will eventually be replaced by the more efficient anisogamy. For the scope of this manuscript we will report whenever the sub-optimal pseudo-isogamic genetic polymorphism can evolve, but we will focus on the evolution of anisogamy assuming it to be the final evolutionary outcome.

By classifying the evolutionary dynamics in this way for all isogamic singular strategy pairs we can then predict the evolutionary dynamics more generally. To do this we first choose the values of the model parameters, and we then numerically solve for the singular strategies (Appendix 5.1) and then get the *attractiveness* and *invadability* (Appendix 5.2) for both the isogamic constrained and unconstrained evolution. We report if these attractors are points where evolution either (1) stops resulting in stable isogamy, (2) transitions into anisogamy, (3) transitions into either anisogamy or pseudo-isogamic genetic polymorphism. We iterate this procedure for a wide range scenarios by systematically varying the parameters of the model.

A special case occurs if there is a single attractor of the isogamic constrained evolution 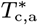 and either all repellers 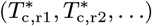 are on the edge of the trait space (occurs for minimum gamete size 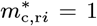, no competition investment 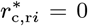, or full competition investment 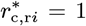) or alternatively if there are no repellers. Then 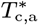 is an unequivocal destination of the isogamic evolution. In all other cases, there might be multiple evolutionary outcomes, and which one is reached might depend on the initial trait value from where evolution starts and the mutational covariance matrix C.

If anisogamy evolves (cases (2) and (3)) the isogamic constraint is removed and the number of evolving traits is increased from two: *T_c_* = (*m_c_, r_c_*), to four: *T* = (*m_x_, r_x_, m_y_, r_y_*). We can numerically solve for the isogamic singular strategies 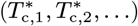, but we can no longer solve for the singular strategies of the unconstrained evolution 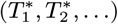 as the dimension of their trait space is too high. To get the evolutionary outcome after anisogamy evolves we have to rely on simulation of the evolutionary path.

### Simulating the evolutionary path

In parallel with analysing the isogamic singular strategies predicting the evolutionary outcomes, we also numerically simulate the evolutionary path given by Eq. 6 (as described in Appendix 5.3). We start the simulation of the evolutionary path under the isogamic constraint and let it run until it stabilises at an isogamic attractor 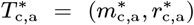. We can then verify if the simulated result matches the predictions from the analysis (presented above) of the *attractiveness* and the trait value of the singular strategies. Then, we continue the numerical simulation without the isogamic constraint, starting at the corresponding singular point of the unconstrained evolution 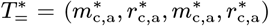 allowing for anisogamy to evolve (ignoring the possibility for isogamic polymorphism to evolve in case (3) as mentioned above). If 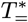 is an attractor no evolution follows, the population stays isogamic and we conclude that 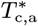 was an isogamic end-point of evolution. On the other hand, if 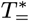 is a repeller, trait values for the different mating types diverge away and anisogamy evolves. We can here verify again that the simulated result matches the predictions gained from the stability analysis of the singular strategies of whether anisogamy evolves or not.

Finally, we let the simulation of the unconstrained evolution run until the evolutionary path stabilises at an anisogamic attractor 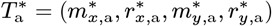 giving us the anisogamic end-point of evolution (a fixed point attractor), or until evolution stabilise in an oscillating cycle (a limit cycle attractor). Since we could not solve for the anisogamic singular strategies, simulations give additional results that could not be obtained with the stability analysis of singular point presented above.

### Parameter reduction and gamete density

We find that the number of parameter can be reduced down to five when analysing the evolutionary dynamics of the model. The model has a total of nine parameters (*d*, *e*, *K_g_*, *K_z_*, *p*, *t*_end_, *v*_0_, *α_m_* and *α_r_*) and we found that five of the parameters always occur as a product together in the invasion fitness function (Eq. 4) and thereby also in the the selection gradient (Eq. 5) as well as in the evolutionary path (Eq. 6). These parameters are *d*, *e*, *p*, *t*_end_ and *v*_0_. Hence, when analysing the evolutionary dynamics we can substitute their product into a single parameter *δ* = *d* × *e* × *p* × *t*_end_ × *v*_0_ reducing the numbers of free parameters of the models down to a total of five: *δ*, *K_g_*, *K_z_*, *α_m_* and *α_r_*.

Varying the parameter *δ* is the same as varying one or more of the five parameters of its product (*d*, *e*, *p*, *t*_end_ and *v*_0_). Hence, varying *δ* can correspond to varying many different properties of the system. First, varying *δ* can correspond to varying the gamete density, as it contains *d* × *e*: the two parameters being proportional to the initial gamete density *n*_*i*, 0_. This means that, among many things, the value of *δ* regulates whether the system tends towards *gamete limitation* at low gamete densities or *gamete competition* at high gamete densities. Furthermore, *δ* contains *t*_end_ × *v*_0_ which is proportional to the distance travelled per gamete, and also *p* the probability of a collision resulting in fertilisation. All five parameters encapsulated in *δ* have identical effects on the evolutionary dynamics (as they occur together in a product): they regulate whether the system tends towards *gamete limitation* low values of *δ*, or *gamete competition* high values of *δ*. From now on, for the sake of simplicity we will refer to *δ* as gamete density (*d × e*), assuming the other parameters (*p*, *t*_end_, and *v*_0_) to be constant. Note that when we vary the gamete density *δ* = *d* × *e*, it can both correspond to changing from low to high population density *d*, or from low to high amount of energy allocated to gamete production per individual *e*.

## Results

Here, we present how two gametic traits, gamete size *m_i_* and relative competition investment *r_i_*, coevolve in a population with two mating types *x* and *y*, where the strategy of a genotype is given by the four gamete traits *T* = (*m_x_,r_x_,m_y_,r_y_*). We begin with some general results of the stability analysis for the transition from isogamy to anisogamy. We then present the simulation results for the coevolution of gamete size and sex-bias in competition in two mating types. We perform the analysis for the parameter range presented in Figures S1–S2 representing a a wide range of scenarios (with subsets presented in Figures 4–5).

### Transition from isogamy to anisogamy

We find that for any set of parameters within our investigated parameter ranges (Figures 4–5 and S1–S2) there is always a single isogamic strategy to which the population converges. This occurs because among the singular strategies of the isogamic constrained evolution there is always a single attractor *T*_c,a_ = (*m*_c,a_, *r*_c,a_) and the rest (one or more) are repellers on the edge of the trait space (*m*_c_ = 1, *r*_c_ = 0, or *r*_c_ = 1) for each parameter combination. Consequently, evolution will always approach the attractor *T*_c,a_ independently of from where it starts. After the isogamic constrained evolution reaches this attracting strategy *T*_c,a_, we find that the evolution either stops and the population stays isogamic (e.g. Figure 3a) or anisogamy evolves as selection turns disruptive for the gamete traits of the two mating types. This results in divergence between two mating types *x* and *y* of both their gamete sizes (*m_x_* and *m_y_*) and their relative competition investments (*r_x_* and *r_y_*) (Figure 3b-d). For both types of competition traits, we show in Figures. 4–5, S1–S2 the parameter spaces where evolution leads to isogamy (within dashed contour lines) or anisogamy (outside dashed contour lines). This shows that anisogamy is generally the more common outcome at low gamete densities *δ* and at high survival constraint on either gametes *K_g_* or zygotes *K_z_*, regardless of the competition trait that gamete size coevolves with.

**Fig. 3.**
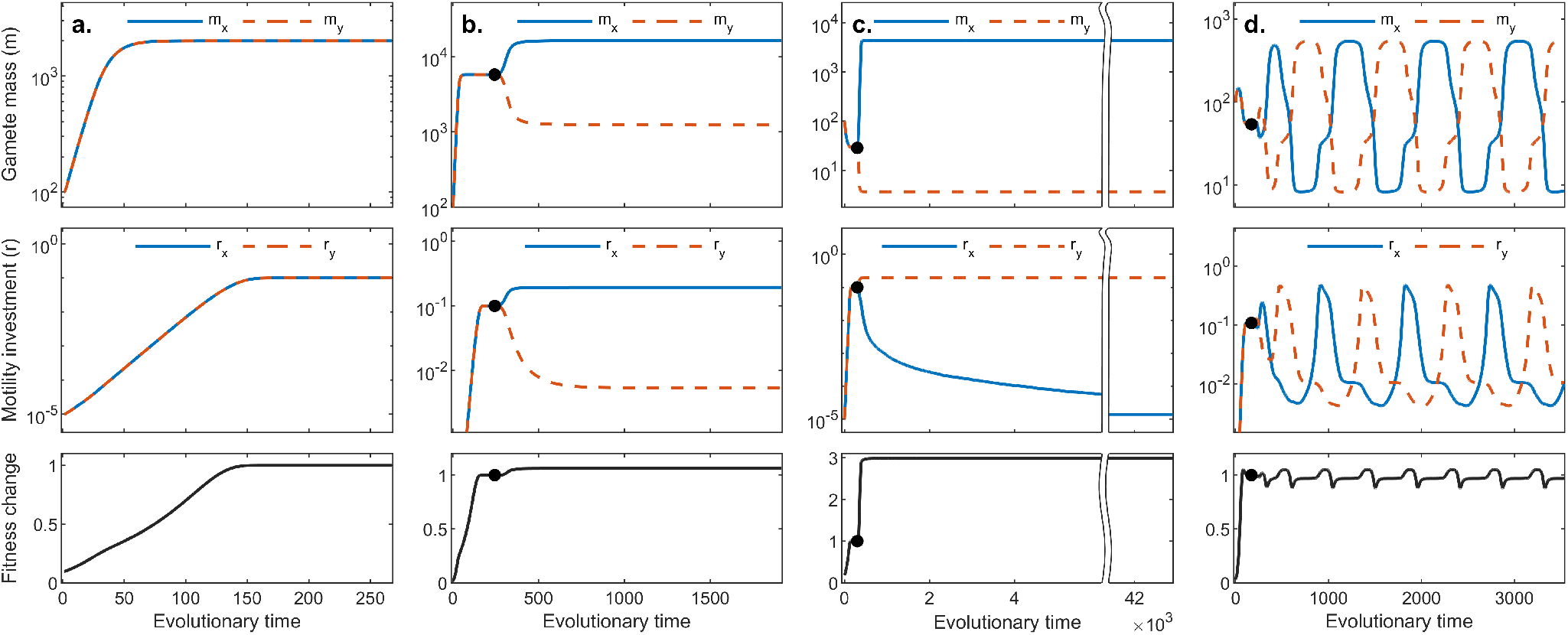
Four examples of simulation runs depicting qualitatively all possible evolutionary outcomes of the model: **a.** no evolution of sexual dimorphism in gamete size or competition, **b.** evolution of anisogamy with female-biased competition investment, **c.** evolution of anisogamy with male-biased competition investment and **d.** oscillating evolutionary dynamics (limit cycle) alternating between female-biased and male-biased competition investment. In each graph, the horizontal axis represents evolutionary time in equation 6, and the vertical axis gives trait value for gamete mass (top panels), competition investment (middle panel) and how fitness changes (Eq. A9) relative to the isogamic attractor marked out by a black dot (bottom panels). Values are given for the two mating types *x* (blue, full line) and *y* (orange, dashed line). If stable anisogamy evolves (cases **b.** and **c.**), the mating type with the larger gamete size is called female and denoted *x* (blue) while the one with smaller gametes is called male and denoted *y* (red). Parameters for figure **a.**-**c.**: *K_g_* = 10^4^, *K_z_* = 0 and *δ* = 0.01, *α_r_* = 0.2 and *α_m_* = (0, −0.4, 0.4) respectively for **a.,b.** and **c** (parameter combinations indicated by stars in figures 4 and S1). Parameters for figure **d.**: *K_g_* = 30, *K_z_* = 10^4^ and *δ* = 10, *α_r_* = 0.6 and *α_m_* = −0.4 (parameter combination indicated by a star in figure S1).

**Fig. 4.**
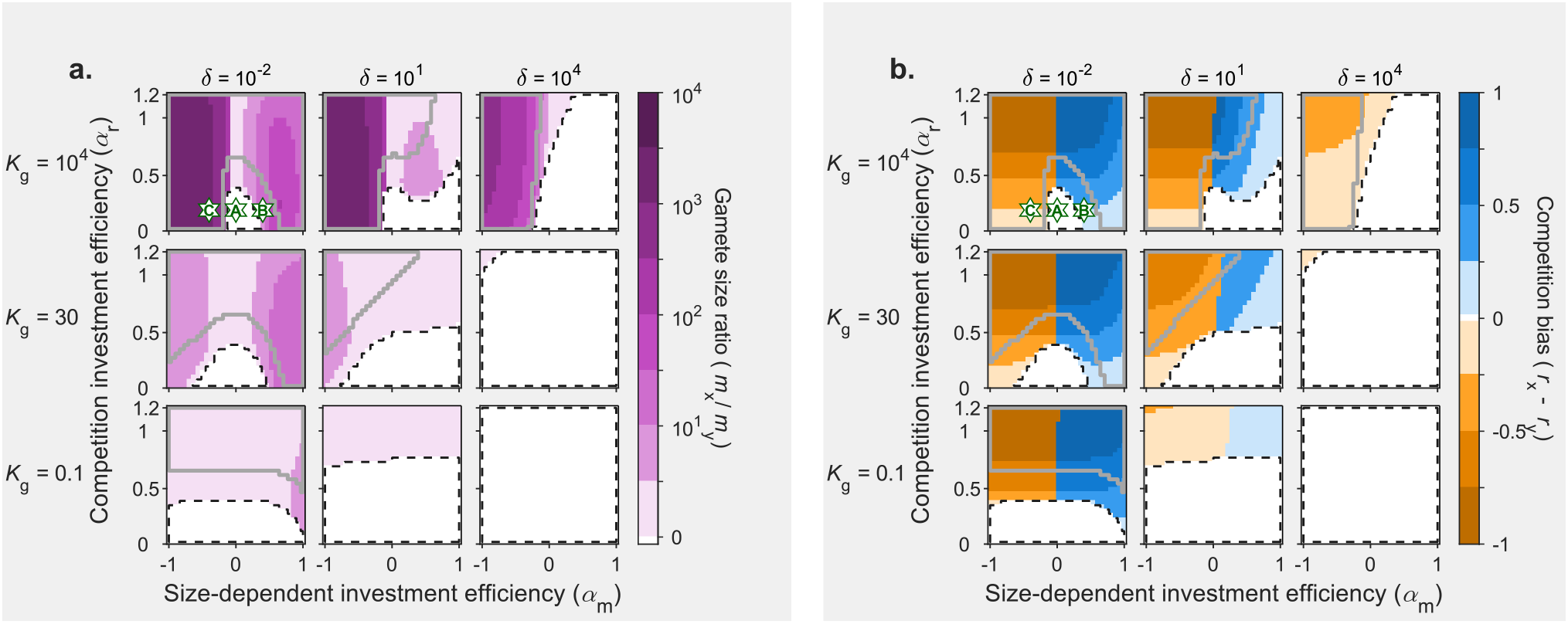
Sex-bias in motility (**b.**) and sexual dimorphism in gamete size (**a.**) at the evolutionary endpoint, as a function of size-dependent investment efficiency *α_m_* (horizontal axis of each subplot) and competition investment efficiency *α_r_* (vertical axis). Size-dependent investment efficiency *α_m_* modulates whether gamete size has a positive or negative effect on the efficiency of energy invested in the competition trait (*α_m_* > 0 or *α_m_* < 0, respectively). The higher the value of competition investment efficiency *α_r_*, the more profitable it is to invest more energy into motility. Graphs are shown for three values of gamete density *δ* (columns), which represent variation from a *gamete limitation* context (*δ* = 0.01) to a *gamete competition* context (*δ* = 10 000), and for three values of the gamete survival constraint *K_g_* (rows). For all graphs, zygote survival constraint *K_z_* = 0. Contour lines correspond to derivations of the stability analysis (identical for **a.** and **b.**). A black dashed line encapsulates the area where isogamy is the expected evolutionary end-point; in the remaining area anisogamy is expected, and a grey contour encapsulates the area where a pseudo-isogamic genetic polymorphism can occur before anisogamy evolves. Coloured shading gives the results from the numerical simulations. Colour intensity expresses the degree of sexual dimorphism in gamete size (**a**) or investment in competition (**b**) at the evolutionary endpoint. White represents no dimorphism and deep colours represent strong dimorphism. Stars represent the parameter combinations for which a simulation run is shown in Figure 3, with the upper-case letter (A,B,C) referring to the corresponding lower-case letter in the panels of Figure 3.

**Fig. 5.**
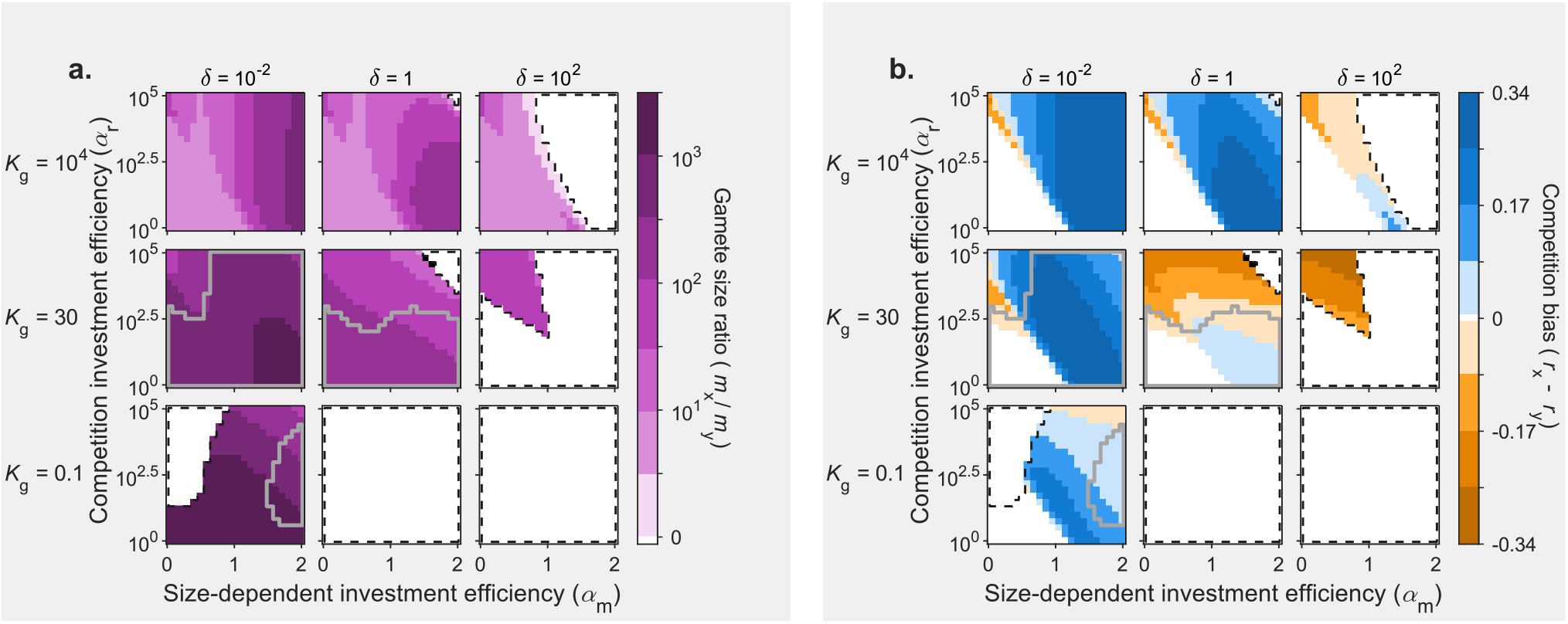
Sex-bias in fusion partner capture (**b.**) and sexual dimorphism in gamete size (**a.**) at the evolutionary endpoint, as a function of size-dependent investment efficiency *α_m_* (horizontal axis of each subplot) and competition investment efficiency *α_r_* (vertical axis). Size-dependent investment efficiency *αm* modulates whether gamete size has a positive or negative effect on the efficiency of energy invested in the competition trait (*α_m_* > 0 or *α_m_* < 0 respectively). The higher the value of competition investment efficiency *α_r_*, the more profitable it is to invest energy into motility. Graphs are shown for three values of gamete density *δ* (columns), which represent variation from a *gamete limitation* context (*δ* = 0.01) to a *gamete competition* context (*δ* = 100), and for three values of the gamete survival constraint *K_g_* (rows). For all graphs, zygote survival constraint *K_z_* = 10^4^. Contour lines correspond to derivations of the stability analysis (identical for **a.** and **b.**). A black dashed line encapsulates the area where isogamy is the expected evolutionary end-point; in the remaining area anisogamy is expected, and a grey contour encapsulates the area where a pseudo-isogamic genetic polymorphism can occur before anisogamy evolves. Coloured shading gives the results from the numerical simulations. Colour intensity expresses the degree of sexual dimorphism in gamete size (**a**) or investment in competition (**b**) at the evolutionary endpoint. White represents no dimorphism and deep colours represent strong dimorphism.

For a considerable subset of the parameter region where the evolution of anisogamy is expected, there is the possibility for the suboptimal pseudo-isogamic genetic polymorphism to evolve (outlined with grey contours in Figures 4–5 and S1–S2). In that case, evolution can result in two diverging isogamic genotypes, one producing large gametes (of both mating types) and the other producing small gametes. However, as soon as the isogamic constraint is removed this situation is no longer stable and anisogamy evolves. We note that in these cases, anisogamy is probably the most likely outcome of evolution, as it is up to twice as efficient (see Analysis section). For this reason, we assume in the simulations that anisogamy evolves under these conditions.

### Simulation of the coevolution of gamete size and competition in two mating types

Solving for the isogamic singular strategies and analysing their evolutionary stability told us whether anisogamy will evolve or not. In addition, we also simulate the evolutionary path (Eq. 6) with the double aim of confirming the result of the stability analysis and finding the trait values at the anisogamic endpoint of evolution *T* = (*m_x_*,*r_x_,m_y_,r_y_*). Simulations do not include the possibility for pseudo-isogamic polymorphisms, directly proceeding to anisogamy instead. Overall, simulations confirm our predictions (with single pixel differences in some places, likely due to numerical imprecisions). For the parameter regions where anisogamy evolves we present simulated results for the degree of gamete size-dimorphism measured as the ratio of gamete sizes of the two mating types *m_x_*/*m_y_*, and the sex-bias in competition measured as the difference *r_x_* - *r_y_* in Figures 4–5, S1–S2. We find that four qualitatively different types of anisogamic outcomes are possible. First, anisogamy can result in either female-bias (*r_x_* > *r_y_*, Figure 3) or male-bias (*r_x_* < *r_y_*, Figure 3c) in competition represented as blue or orange regions, respectively, in Figures 4–5b, S1–S2b). Then, for the fusion partner capture model there are some parameter region where competition trait don’t evolve resulting in anisogamy without bias in competition (*r_x_* = *r_y_* =0, Figure 3c) seen as white regions outside the dashed contours in Figures 5b and S2b). Finally, there are some restricted parameter regions where the anisogamic attractor is a limit cycle resulting in never ending oscillating dynamics (Figure 3d) where male and female bias in competition alternate, represented as black regions in Figures 5, S1–S2). In the next two subsections, we discuss for each competition trait separately the influence of the model parameters on the evolution of anisogamy and sex-bias in competition. For each trait, we vary gamete density *δ* from *gamete limitation* to *gamete competition*, the gamete and zygote survival constraint, *K_g_* and *K_z_* as well as competition investment efficiency *α_r_* and size-dependent investment efficiency *α_m_*.

#### Competition through gamete motility

Figure 4 presents a subset of the results with zygote survival constraints absent (*K_z_* = 0, all zygotes formed survive) for simplicity as it has only a minor effect. We can see that anisogamy can evolve under a wide range of gamete densities *δ*, provided that there is a strong constraint on gamete survival (large *K_g_*, Figure 4a, top row). If the constraint on gamete survival is somewhat relaxed (Figure 4a rows two and three), anisogamy may only evolve under lower gamete densities. This result is reminiscent of the classical finding that a constraint on gamete survival encourages the evolution of anisogamy in the *gamete competition* context (Bulmer and Parker, 2002). A comparison between Figure 4a and b shows that the evolution of anisogamy for this competition trait is always connected with the evolution of a sex-bias in motility.

Each subplot of Figure 4 also depicts the effects of competition investment efficiency *α_r_* and size-dependent investment efficiency *α_m_*, the two parameters that modulate the efficiency of investing energy into motility. First, the competition investment efficiency *α_r_* either facilitates the evolution of anisogamy or has no effect. Furthermore, *α_r_* increases the sex-bias in competition trait whenever anisogamy evolves. Looking at Figure 4b, we can see that the size-dependent investment efficiency *α_m_* dictates which sex invests more energy into motility: when *α_m_* is positive, large gametes profit more from investing into motility and in that region of the parameter space we observe exclusively female-biased investment in competition if any sex-bias is present. Inversely, when *α_m_* is negative, indicating that smaller gametes benefit more from investing into motility, we observe that only a male-bias in competition is possible. We can further note that this situation is not symmetric, at least at higher gamete densities (Figure 4b columns two and three): a male-bias in competition appears more easily (i.e. for lower values of *α_r_*) than a female-bias in competition. At the highest gamete density, a female-bias in competition is not observed at all, and anisogamy may only evolve if smaller gametes benefit more from investing in motility. However, for the lowest gamete density (Figure 4b column 1), the effect of *α_m_* is rather symmetric and it is not clear whether the sex-bias in competition appears more easily in males or in females (it appears to vary slightly with *K_g_*). Another trend worth noting is the fact that when anisogamy evolves with a male-bias in competition, gamete size difference is of larger magnitude than when it evolves along with a female-bias in competition. This intuitively makes sense because investment in competition will divert energy away from gamete mass to competition: when males spend more on competition than females, this will magnify gamete size difference by making the small gametes smaller, and *vice versa*. Both *K_z_*, the zygote survival constraint and *K_g_*, the gamete survival, have a positive effect on the evolution of anisogamy in high gamete densities (see Supplementary Figure S1).

We want to highlight three main points from this section of the results: (i) when anisogamy evolves it seems to almost always be accompanied by a sex-bias in competition, (ii) the nature of the competition trait (modulated by *α_r_* and *α_m_*) influences the possibility for the evolution of anisogamy as well as the direction of the resulting sex-bias in competition, and finally (iii) whereas only male-biased competition can evolve under high gamete density (*gamete competition*), both male and female bias in competition can evolve under low density (*gamete limitation*).

#### Competition through fusion partner capture

Here, we investigate the coevolution of gamete size with a second competition trait which increases apparent gamete size in order to capture nearby gametes to achieve fertilisation. An increase in apparent size will improve the target size of the gamete but will not increase survival probability. Similarly to the scenario with motility as a competition trait, we find that the evolution of anisogamy is mostly paralleled by the evolution of a sex-bias in competition (Figures 5,S2). A noticeable difference however, is that when both *α_m_* and *α_r_* are small it is frequent that none of the mating types invest in the competition trait, because it is not energy-efficient enough (see for example lower left corner of each subplot in Figure 5b). As in the motility scenario, high survival constraints on gametes and zygotes (high values of *K_g_* and *K_z_*) also facilitate the evolution of anisogamy, with the difference that with this competition trait a higher overall constraint seems required for anisogamy to evolve; for this reason, we show a subset of the parameter space with high zygote survival constraint in Figure 5. A final point of similarity between the two competition scenarios (motility and fusion partner capture) is the fact that a male-bias in competition appears more likely at high gamete density (*gamete competition*) than at low density (*gamete limitation*). There are also striking differences between the motility and fusion partner capture scenarios: when fusion partner capture is considered as a competition trait, the possibility for anisogamy to evolve at high gamete density is more restricted than in the motility case. Also, at lower gamete densities, anisogamy is in most cases accompanied by a strong female-bias in competition and male-bias competition is almost absent. This again shows the great influence of the competition trait considered for the possibility for anisogamy to evolve and the resulting sex-bias in competition. The influence of *α_r_* and *α_m_* on the evolution of anisogamy and the sex-bias in competition appears less straightforward than in the motility scenario. Increasing *α_r_* sometimes facilitates and sometimes prevents the evolution of anisogamy. Whenever male-bias and female-bias co-exist within the same combination of parameters *K_g_*, *K_z_* and *d* (Supplementary Figure S2c and e, top rows), the transition from one to the other isn’t defined by a single pivot value of *α_m_*, as was the case for the motility scenario. Instead, the transition from female-bias to male-bias appears to be a function of *α_m_*, *α_r_* and survival constraints.

We have seen that contrary to the motility trait, fusion partner capture evolves overall more easily in larger gametes. This result may appear obvious, because the cost/benefit relationship for this trait is by construction in favour of larger gametes that can expand their apparent size at a limited survival cost. However, we advocate that this result is still meaningful, in showcasing the variety of competition trait that may arise by chance, some more efficient in large gametes and some in small gametes. Examining this diversity is necessary to fully understand the relationship between the evolution of anisogamy and that of intrasexual competition. On the contrary, making the restrictive assumption that competition should generally favour small gametes (by only considering motility with negative *α_m_* for example) can lead to the conclusion that males should invest more into competition, but this view leaves many possible scenarios aside. With this second competition trait, we show that the evolution of anisogamy and sex-bias in competition is also possible when competition happens through a trait favouring female strategies (large gamete). We elaborate more on this point in the discussion.

## Discussion

### Two pathways to anisogamy and resulting sex-bias in competition for fertilisation

We have presented a mechanistic model of the coevolution of gamete size and a competition trait at the gamete level that increases individual reproductive success. Our work shows that anisogamy can evolve along with a sex-bias in intrasexual competition. This is true for the two different scenarios of competition traits investigated here, first a gamete motility competition trait and second a fusion partner capture competition trait. Previous work has shown that anisogamy may have evolved under *gamete competition* (high gamete density) or *gamete limitation* (low gamete density) (e.g. Bulmer and Parker, 2002; Lehtonen and Kokko, 2011; Hoekstra et al., 1984). We believe that this distinction is important because the forces of selection that would have acted to produce gamete size differences in these two scenarios are not the same. This has implications for the evolution of intrasexual competition, which we investigate here. This question is relevant to the initial evolution of anisogamy from an isogamous state, but also to the maintenance and continuous evolution of gamete size in contemporary species as we will discuss later.

In the *gamete competition* context, under high gamete density, the forces that are expected to drive the evolution of anisogamy are size-dependent survival selection and competition for fertilisation success. In that context, anisogamy only evolves under a size constraint on survival. If anisogamy does evolve, under high gamete density, all large gametes become fertilised but smaller ones are in excess, leading to the evolution of male-biased investment in intrasexual competition. In the *gamete limitation* context, under low gamete density, the main force expected to drive the evolution of anisogamy is a disruptive selection pressure on gamete size that maximises encounter rates of gametes from the two mating types. In that context, both sexes could be expected to engage in competition because they may both have unfertilised gametes at the end of a mating event. By adjusting gamete density, our model allowed the exploration of *gamete competition* and *gamete limitation* scenarios. We have seen again that anisogamy may evolve in both cases, but with different outcomes in terms of the sex-bias in competition. A *gamete competition* context clearly favoured the evolution of a male-bias in competition, regardless of the competition trait considered. But the evolution of anisogamy was more restricted in that context, relying on a high level of size-dependent survival constraint on gamete or zygote (as in Bulmer and Parker, 2002). On the other hand, a *gamete limitation* context allowed both sexes to invest in competition, with the final sex-bias determined by the nature of the competition trait, more specifically whether it was defined as more energy-efficient in smaller or larger gametes. This result makes evident that the evolution of anisogamy needs not result in the evolution of a male-bias in competition, but may lead to a variety of outcomes depending on the circumstances under which anisogamy initially evolved (*gamete limitation* or *gamete competition*) and which competition trait is considered.

### Competition traits in the model and in nature

We implemented in our model two possible competition traits. Although those traits are gamete-level traits, we remind the reader that our model is focusing on individual-level selection and not gamete-level selection. Indeed, the competition traits represent competition investment at the individual level, as they only influence individual-level reproductive success. Both traits are flexible with the addition of two parameters that allow to describe freely how energy investment and gamete size relate to the efficiency of the trait. This allows us to avoid *a priori* assumptions regarding which sex should benefit more from investing in a given trait. We have shown that the nature of the efficiency relationships of the traits greatly influences the resulting sex-bias in selection.

With the first trait, gamete motility, we have seen that it evolves in males in the *gamete competition* context and can evolve in either sex in the *gamete limitation context*, depending on the relationship between gamete size and trait efficiency (*α_m_*). In short, if small gametes benefit more from investing in that trait then motility will be male-biased and *vice versa*. The fact that few swimming eggs have been observed in nature so far (Motomura and Sakai, 1988; Klochkova et al., 2019) suggests that small gametes may generally swim more efficiently than large ones. We note however that a positive relationship between size and speed of gamete has been reported in a unicellular algae (Seed and Tomkins, 2018), but this relationship may not hold for size differences of a thousand fold or more as is often the case in anisogamous systems. Regardless, it is easy to imagine other traits that could be less costly to develop for larger gametes. With that in mind, we incorporated in our model a second trait, fusion partner capture. This trait increases the apparent size of the gamete, making it a larger fertilisation target without increasing its survival probability. In our model, this trait develops mainly in the mating type producing the larger gamete, resulting in female-biased competition. This trait can be compared to the egg jelly coat found in several species of broadcast spawning marine invertebrates (e.g. Podolsky, 2002; Farley and Levitan, 2001), and seems therefore quite realistic. It it also possible that the trait we model describes somewhat accurately the release of chemoattractant by gametes, which is another way of increasing apparent size. This strategy is found in the eggs of several species, both external and internal fertilisers, including mammals (Eisenbach and Giojalas, 2006).

### Prevailing forces in the evolution and maintenance of anisogamy in animals

In our model, the evolution of anisogamy was almost always accompanied by a sex-bias in competition. *Gamete competition* and *gamete limitation* shaped very different sex-biases in competition. This means that an understanding of the evolutionary pathways that initiated anisogamy is key in the relationship between this initial anisogamy and any resulting sex-bias in competition. The current theory on the evolution of anisogamy in animals proposes that it likely originated in sessile broadcast spawners (Parker, 2014), a type of animals that are often subjected to *gamete limitation* (Levitan, 1996). This then suggests that *gamete limitation* may have been important in the evolution of an initial sexual dimorphism in gamete size. According to our results, it seems therefore unlikely that the evolution of anisogamy should have resulted in male-biased intrasexual competition in all anisogamous animals. Indeed, in our model the *gamete limitation* scenario can lead to either female or male-biased competition, depending on the nature of the competition trait considered.

Whether the sex-bias in competition at the initial anisogamy has an influence on the patterns of sex-specific competition in contemporary anisogamous species is unclear. The complex patterns of sex-specific selection that we observe today may be subjected to an array of confounding factors that we do not account for, such as evolutionary history or ecological constraints. Nevertheless, we want to comment, with our findings in perspective, the fact that in the majority of natural systems studied today more intense intrasexual competition is found in males than in females (Janicke et al., 2016; Janicke and Morrow, 2018). First, the claim of a general male-bias in competition should be nuanced: although males experience more intense intrasexual competition in a majority of species (Janicke et al., 2016; Janicke and Morrow, 2018), there is a lot of variation among taxa as clearly visible from Figure 1 in Janicke et al. (2016). Females do experience non-negligible levels of intrasexual competition in most species and in some cases more than males (reviewed in Hare and Simmons, 2018). Female competition is favoured if males provide costly nuptial gifts or parental care, but they may also compete to increase fertilisation success in sperm limited-contexts (Hare and Simmons, 2018), which are not restricted to external fertilisers (see for example a study in gorillas Niemeyer and Anderson, 1983, and one in saiga antelopes Milner-Gulland et al., 2003). Second, we propose two non-mutually exclusive hypotheses that may explain the apparent inconsistency between our theoretical claim about the origin of anisogamy and empirical observations of today’s natural systems. (i): a large proportion of animals studied by biologists are internal fertilisers. The evolution of internal fertilisation reduces the chances for *gamete limitation* to appear, which in turn should favour the evolution of male competition. (ii): our model assumes that individuals invest the same amount of energy into reproduction regardless of their sex or mating type. If that assumption does not hold, and if the cost of producing larger gametes does not scale linearly with gamete size, it is possible that a difference in potential reproductive rate (PRR, time required for an individual to produce offspring and return to the mating pool, as in Clutton-Brock, Clutton-Brock and Parker, 1992) arises between the sexes. A sex-bias in PRR is likely to cause a sex-bias in intrasexual competition, with the sex with the highest PRR competing for mating opportunities with the other sex (Clutton-Brock and Parker, 1992; Kokko et al., 2006). We note that these two hypotheses (i) and (ii) do not undermine the fact that anisogamy (gamete size dimorphism) is not likely to explain directly sex-biases in intrasexual competition: in the first case it is the evolution of internal fertilisation that causes males to compete more than females, in the second case it is a sex-bias in PRR, which could arise from anisogamy as well as from other causes. For example, PRR is highly sensitive to ecological factors, and field studies have shown that sex-biases in PRR can switch within the course of single mating season (Almada et al., 1995; Forsgren et al., 2004, reviewed in Ahnesjö et al., 2008; Kvarnemo and Ahnesjo, 1996).

If it seems likely that *gamete limitation* would have played an important role in the evolution of anisogamy in ancestral animals, we can also wonder what are the evolutionary forces that maintain anisogamy in contemporary species. We have suggested above that the evolution of internal fertilisation should have created a *gamete competition* context favourable to male competition. This claim should, however, be taken with care. A recent study in mammals has shown an inverse relationship between body size and sperm cell size (Lüpold and Fitzpatrick, 2015), a trend suggesting that in larger mammals *gamete limitation* may happen to some extent, leading to the evolution of smaller more numerous sperm cells, less competitive but increasing chances of fertilisation in low gamete density. Sperm limitation, or failure of females to get all of their eggs fertilised may also be more common than expected in insects (reviewed in García-González, 2004). Finally, many contemporary invertebrate marine species are broadcast spawners that are often subjected to *gamete limitation* (Levitan and Petersen, 1995). In these species, variance in reproductive success may become higher in either sex depending on gamete density (Levitan, 2004), indicating that intrasexual competition may easily arise in either sex, as suggested by our results. Furthermore, a wealth of laboratory and field studies in broadcast spawners have reported female traits that increase fertilisation success and are therefore involved in female intrasexual competition. In three species of sea urchins (Levitan, 1993, 1998a) egg traits are shown to evolve to maximise fertilisation success. In the sea urchin *Lytechinus variegatus* experimental removal of a jelly coat around the eggs lowers fertilisation success due to reduced target size (Farley and Levitan, 2001). In the sand dollar *Dandraster excentricus* a jelly coat that increases up to sixfold the size of the egg increases fertilisation success (Podolsky, 2002), and finally in the tunicate *Styela plicata* (Crean and Marshall, 2008) both sexes adapt their gametes to population density, with female gametes becoming notably larger at low density, a strategy that increases fertilisation success but comes at a cost for zygote survival.

### Conclusion

It seems that *gamete limitation* may be important in shaping anisogamy, which means that the evolutionary forces that are responsible for gamete size differences cannot be expected to always result in the evolution of male-biased competition. We thus question the claim that anisogamy necessarily leads to male-biased competition. Some of the evolutionary forces that can spur the evolution of anisogamy in the *gamete competition* context do clearly favour male-biased competition; but evolutionary forces that shape anisogamy in the *gamete limitation* context may produce male-biased or female-biased competition, with the outcome mostly decided by the nature of the competition trait and how its efficiency relates to gamete size. There are good reason to think that *gamete limitation* played an important role in the animals that first evolved the male and female sex. Even though the evolution of internal fertilisation in a wide range of species suggests that *gamete competition* is common in nature, there is evidence claiming that even in internal fertilisers the evolutionary forces of *gamete limitation* are still at play. Together, this suggests that, as far as anisogamy is concerned, females should benefit from competing for increased fertilisation success, too, a claim that is supported by the observation of female intrasexual competition in many species, an in particular with egg competition traits in broadcast spawning species. Evidence showing more intense competition in males in a majority of species (a general pattern but not a general rule) is not proof that anisogamy is the cause for that pattern. We suggest that, for example, potential reproductive rate (PRR) may be a better predictor of intrasexual competition, as it captures important ecological influences on competition. In turn, PRR may be related to gamete size in some species, if larger gametes require more time and energy to be produced but this does not need to be the case. Our results challenge the classical view that anisogamy alone is enough to explain more intense intrasexual competition in the male sex.

## Appendix

Note: In the main part we present how the variables of the model are functions of the gamete traits (*m_x_*, *r_x_*, *m_y_* and *r_y_*). We will not detail this in the appendix for the sake of readability. One exception is made for the case of fertilisation rates *f_i_* and 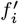 (Eqs. A9 and A14) that are used to obtain the expected evolutionary path of these gamete traits (Eqs. 4, 5, 6 in the main text).

### 1. Collision frequency

To mechanistically model the fertilisation dynamics between the gametes of the two mating types *x* and *y* in the mating pool, we make use of classical collision theory from chemistry and physics (McNaught et al., 2014). Hence, we assume that gametes of the two mating types *x* and *y* travel at constant speeds *v_x_* and *v_y_* with trajectories approximated by straight lines at a local scale. We obtain the following frequency of collisions between gametes of the two mating types (per unit of time and per unit of volume)

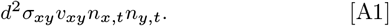

Here, *n_i,t_* is the number of gametes per zygote of mating type *i* at time *t*, and *d* is the zygote population density (number of zygotes per mating type per unit of volume). Also, 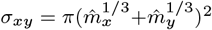 is the area of trajectories which would result in a collision between gametes of the two mating types, representing a disk with a radius equal to the combined radiuses of the two gametes (i.e, their collisional cross section), assuming spherical gametes and excluding the motility machinery from the collision target. Lastly, 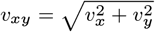 is the average relative velocity between gametes of the two mating types.

### 2. Gamete fertilisation dynamics

Gametes of the two mating types *x* and *y* collide at a frequency given by Eq. A1 and each collision has a probability *p* of resulting in a fertilisation event. Fertilisation generates a juvenile zygote and removes the colliding gametes from the mating pool.

Multiplying Eq. A1 with *p* gives the fertilisation rate (per unit of time and per unit of volume). Dividing the fertilisation rate by the gamete density of mating type *i*, *dn_i,t_*, gives the fertilisation rate per gamete of mating type *i* per unit of time at time *t*. Multiplying this rate with the number of gametes per zygote of mating type *i*, *n_i,t_*, gives the fertilisation rate per zygote of mating type *i* per unit of time at time *t*.

Because gametes that achieve fertilisation are removed from the mating pool, the number of gametes present at time *t* per zygote *n_i,t_* decreases according to their fertilisation rate (negative term on the right hand side of the equation)

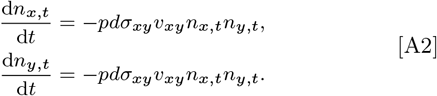

The number of gametes per zygote of the two mating types declines at the same rate (d*n_x,t_*/d*t* = d*n_y,t_*/d*t* in Eq. A2) as each fertilisation always removes one gamete of each mating type. The difference in the number of gametes between the two mating types *n_y,t_* – *n_x,t_* thereby stays constant over the mating period and equals the initial difference *n*_*y*, 0_ – *n*_*x*, 0_. The initial number of gametes per zygote of mating type *i* equals

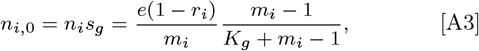

where *n_i_* is the number of gametes produced per zygote of mating type *i* and *s_g_* the survival probability of each gamete (see main text). Hence, the number of gametes per zygote of one mating type at time *t* is given by the number of gametes of the other mating type plus their initial difference

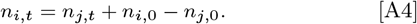

where *i* is the focal mating type (*x* or *y*) and *j* is the other mating type.

By using the substitution in Eq. A4 we can express equation A2 as a single differential equation (as the initial gamete densities *n*_*i*, 0_ and *n*_*j*, 0_, Eq. A3, are constants)

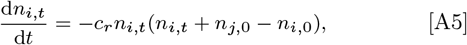

where *c_r_* = *pdσ_xy_v_xy_* is a constant proportional to the fertilisation rate. Eq. A5 is a separable differential equation with the following solution

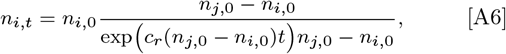

(step-wise calculations in Supplementary material 1.1, Eq. S3).

#### 2.1. Gamete fertilisation dynamics under isogamy

For the case of evolution under isogamic constraint, the gametes of the two mating types are constrained to equal trait values. Hence, we have that *n_x,t_* = *n_y,t_* and the change in the number of gametes over time Eq. A5 simplifies to

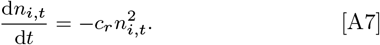

Solving for *n_i,t_* in Eq. A7 gives the number of gametes per zygote of mating type *i* at time *t* under the isogamic constraint

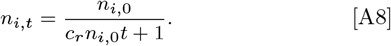

(step-wise calculations in Supplementary material 1.2, Eq. S6).

### 3. Zygote fertilisation success

When the mating period is over, the number of fertilised gametes per zygote equals *n*_*i*, 0_ − *n*_*i,t*_end__ (Eqs. A3 and A6), (i.e., the number of gametes entering the mating pool subtracted by the number of unfertilised gametes at *t* = *t*_end_, the end of the mating period). Each fertilisation produces a juvenile zygote (Fig. 2) that has a probability of survival to the next generation of *s_z_* = (*m_x_* + *m_y_* – 1)/(*K_z_* + *m_x_* + *m_y_* – 1) (as presented in the main text). Hence, the number of fertilised gametes per zygote multiplied with their survival probability *s_z_* gives the per capita fertilisation success *f_i_* (or simply: fertilisation success) corresponding to the absolute fitness of a zygote,

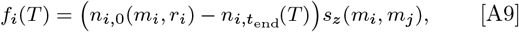

where *i* denotes the focal mating type (*x* or *y*) and *j* the other mating type, and where the trait vector *T* = (*m_x_*, *r_x_*, *m_y_*, *r_y_*) represents all four gamete traits.

### 4. Fertilisation dynamics of a rare mutant strategy

To introduce evolution into our model we assume that mutation events are rare, each introducing a single mutant zygote. The strategy of the mutant 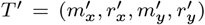 deviates in one or more of the four traits from the resident strategy *T* = (*m_x_, r_x_, m_y_, r_y_*) (if evolution occurs under the isogamic constraint *T*′ is replaced with with 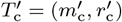 and *T* with *T*_c_ = (*m*_c_, *r*_c_)). The resident population is assumed to be large, and as the mutant strategy *T*′ is introduced at the minimum frequency (one individual) its effect on the fertilisation dynamics can be neglected, as any gamete is very unlikely to collide with these rare mutant gametes. One can thereby consider gametes to only collide with gametes of the resident strategy, as long as the mutant is rare. This simplification is used when deriving the invasion fitness of the a mutant strategy. To obtain the invasion fitness of the mutant (Eq. 4), we need its zygote fertilisation success (Eq. A9), which in turn requires a solution for the gamete fertilisation dynamics of the mutant (Eq. A12), given below.

#### 4.1. Gamete fertilisation dynamics of a rare mutant strategy

Here, we derive the gamete fertilisation dynamics of a rare mutant strategy. The mutant gametes are removed from the mating pool at each successful fertilisation, and the number of gametes per mutant zygote *n_i′, t_* (of mating type *i* at time *t*) decreases according to their fertilisation rate (negative term on the right hand side)

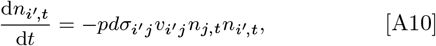

where *n_j,t_* is the number of gametes per resident zygote of the other mating type *j, σ_i′ j_* the collisional cross section between the mutant gamete of mating type *i* and the resident gamete of mating type *j*, and *v_i′j_* is average relative velocity (see Appendix 1) between the mutant gamete of mating type *i* and the resident gamete of mating type *j*.

To solve for *n_i′ t_* we first substitute the number of resident gamete *n_j,t_* in Eq. A10 with Eq. A6 giving

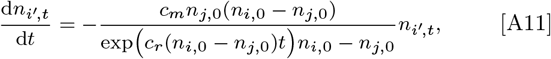

where *c_m_* = *pdσ_i′ j_ v_i′j_* is a constant proportional to the collision probability for the mutant gametes of mating type *i*. Eq. A11 is a separable differential equation with the following solution

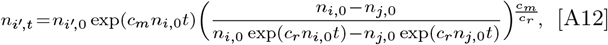

giving the number of unfertilised gametes per mutant zygote at time *t* (step-wise calculations in Supplementary material 1.3, Eq. S11).

If evolution occurs under the isogamic constraint in Eq. A10 we instead substitute the resident gamete number *n_j,t_* with Eq. A8 and we get

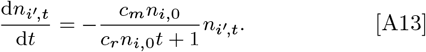

which also is a separable differential equation and has the following solution

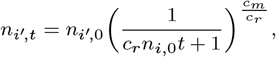

giving the number of unfertilised gametes per mutant zygote at time *t* in an isogamic population (step-wise calculations in Supplementary material 1.4, Eq. S14)

#### 4.2. Zygote fertilisation success of a rare mutant strategy

The fertilisation success of a rare mutant strategy 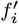 (of mating type *i*) follows the same logic as for the resident fertilisation success (Eq. A9) and is given by

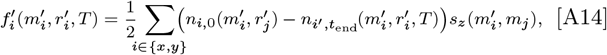

where *i* denotes the focal mating type (*x* or *y*) and *j* the other mating type, and where the trait vector *T* = (*m_x_, r_x_, m_y_, r_y_*) represents all four gamete traits.

### 5. Predicting the evolutionary dynamics

Below we describe our procedure for predicting the evolutionary dynamics of the gamete traits *m_x_*, *r_x_*, *m_y_* and *r_y_* (or *m_c_* and *r_c_* under the isogamic constraint).

#### 5.1. Solving for isogamic singular strategies

We assume evolution to start under the isogamic constraint (where *m_x_* = *m_y_* = *mc* and *r_x_* = *r_y_* = *r*_c_ and the isogamic strategy is given by *T*_c_ = (*m*_c_, *r*_c_)). To predict the evolutionary path (given by Eq. 6) we first numerically solve for the isogamic singular strategies 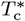, i.e. where the selection gradient is zero, 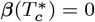. The singular strategies represent candidates for attracting strategies. To find these isogamic singular strategies, we numerically solve for when the first and the second element of *β*(*T_c_*) are equal to zero, separately (i.e. we solve for both isoclines). We then find their intersections, which gives the singular points.

#### 5.2. Stability analysis of singular strategies

To predict the evolutionary dynamics in the vicinity of a singular strategy *T*\* (or 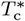 under the isogamic constraint), we look at two stability properties of the singular point, namely the *attractiveness* and *invadability*. First, a singular strategy *T*\* is uninvadable by nearby mutants if the four-dimensional Hessian matrix H of the selection gradient (Eq. 5) with entries

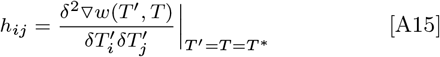

is negative definite (where 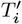 and 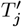 gives the *i*th and the *j*th element of the mutant trait vector 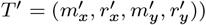. Otherwise, the singular strategy *T*\* is invadable by nearby mutant strategies (Leimar, 2009). We refer to this property as the *invadability*.

Note: if isogamic constraint is considered, replace the gamete strategies *T* and *T*′ with the constraint strategies *T*_c_ and 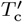, respectively. This gives the two-dimensional Hessian matrix of the constraint evolution. The same holds for the Q-matrix and Jacobian J presented below.

Whether a singular strategy *T*\* is an attractor of the evolutionary path or a repeller depends on the Jacobian matrix J of the selection gradient (Eq. 5). The Jacobian is given by J = H + Q where Q is a four-dimensional square matrix with entries

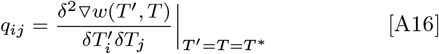

where *T_j_* gives the *j*th element of the resident trait vector *T* = (*m_x_, r_x_, m_y_, r_y_*))

If J is negative definite *T*\* is an attractor meaning that evolution will converge towards *T*\*. On the other hand, if J is positive definite *T*\* is a repeller of the evolutionary dynamics. Lastly, if J is indefinite the evolutionary dynamics will depend on the mutational covariance matrix C. Then *T*\* can be an attractor for specific mutational covariances of C (such that the Jacobian of C*β*(*T*) is negative definite), but is otherwise a repellor of the evolutionary dynamics (see Leimar, 2009).

For our analysis, we do not consider any mutational covariance between our four traits (*m_x_, r_x_, m_y_, r_y_*) (or two traits (*m*_c_, *r*_c_) for evolution under isogamic constraint) corresponding to C being a diagonal matrix, and consequentially *T*\* is only an attractor if J is negative definite, and is otherwise an repeller. For simplicity we choose C to be equal the identity matrix meaning that the evolutionary path Eq. 6 equals the selection gradient Eq. 5.

#### 5.3. Numerical simulation of the evolutionary path

As a complement to the stability analysis, we numerically simulate the evolutionary path given by Eq. 6 using the Runge-Kutta method with adaptive step size of order 4 and 5. First, we simulate the evolutionary path under the isogamic constraint. As we find that there is always a single attracting strategy for the parameter range we investigate (see Results section), the starting point has no particular meaning and we start evolution from an arbitrary point chosen to be *T*_c_ = (*m*_c_ = 100, *r*_c_ = 10^−5^). We run the numerical simulation until the evolutionary equilibrium is reached at an attractor 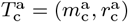. Then, we continue the numerical simulation without the isogamic constraint with a small initial divergence *ε* in either gamete size *m* or relative competition *r* investment traits between the two mating types such that 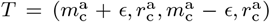 or 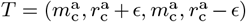. If evolution in both cases leads back towards 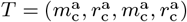 we conclude that the isogamic attractor 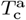 is also an attractor of the corresponding unconstrained evolution. In this case, there is stabilising selection preventing the evolution of anisogamy and the isogamic attractor 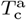 is the end-point of evolution. Otherwise, if the trait values of the two mating types diverge away from 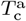, it is a repellor of the unconstrained evolution and anisogamy evolves. We then run the numerical simulation until anisogamic evolution reaches an attractor, most often it is a fixed point attractor 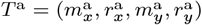 giving an evolutionary endpoint (see Fig. 3b-c), but for a small subset of the parameter values it reaches a limit cycle attractor, resulting in stable oscillations (see Fig. 3d).

## ACKNOWLEDGMENTS

We thank Ingrid Ahnesjö, David Berger, Luc Bussière, Charlotta Kvarnemo, Johanna Liljestrand Rönn and Claus Rueffler for their interest in our project, fruitful scientific discussions and helpful comments to our manuscript.

## Supplementary information

### 1. Step-wise calculations

#### 1.1. Solving for gamete fertilisation

In the mating pool, the number of gametes per zygote of mating type *i* at time *t* decreases as fertilisation events occur according to (Eq. A5)

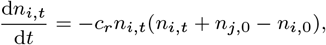

which is a separable differential equation:

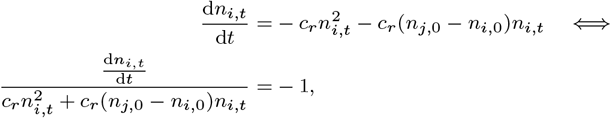

and can thereby be solved by integrating both sides with respect to *t*

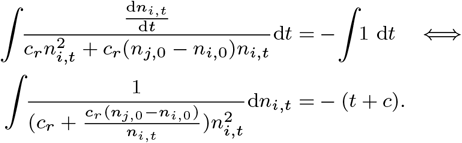

The left-hand side integral can be solved with the following substitution *u* = *c_r_* + *c_r_* (*n*_*j*, 0_ – *n*_*i*, 0_)/*n_i,t_* and 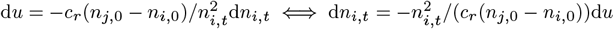, resulting in

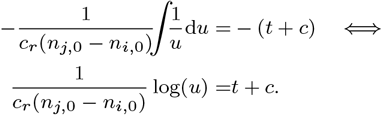

Substituting back *u* = *c_r_* + *c_r_*(*n*_*j*, 0_ – *n*_*i*, 0_)/*n_i,t_* gives

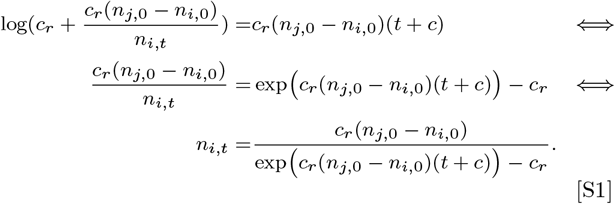

Solving for *c* in Eq. S1 with initial condition *t* = 0 and *n_i,t_* = *n*_*i*, 0_ gives

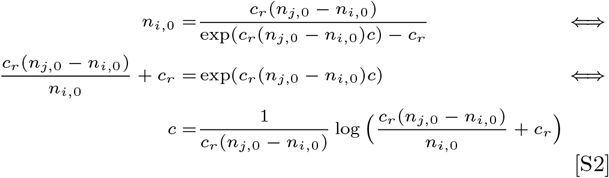

Substituting *c* (Eq. S2) into Eq. S1 gives

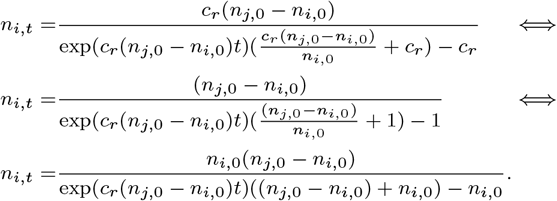

Simplifying this once more gives our explicit expression for the number of gametes of mating type *i* at time *t*

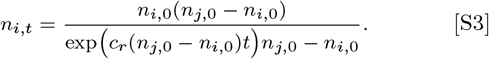

#### 1.2. Solving for gamete fertilisation under isogamy

For the case of the isogamic constraint, the number of gametes per zygote of mating type *i* at time *t* decreases according to (Eq. A7)

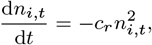

which is a separable differential equation:

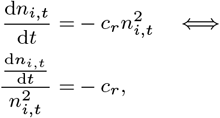

and can thereby be solved by integrating both sides with respect to *t*

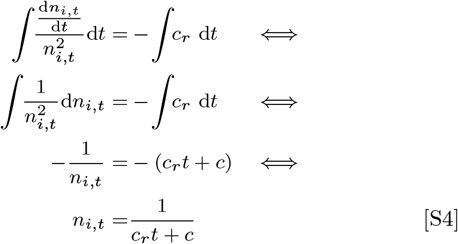

Solving for *c* in Eq. S4 with initial condition *t* = 0 and *n_i,t_* = *n*_*i*, 0_ gives

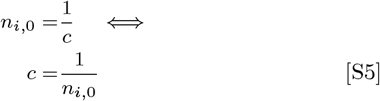

Substituting *c* (Eq. S5) into Eq. S4 gives

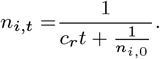

Simplifying this gives our explicit expression for the number of gametes of mating type *i* at time *t* under isogamy

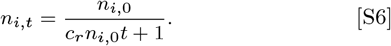

#### 1.3. Solving for gamete fertilisation of a rare mutant strategy

In the mating pool, the number of gametes per zygote with a rare mutant strategy, of mating type *i* at time *t*, decreases according to (Eq. A12)

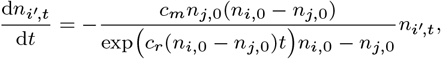

which is a separable differential equation

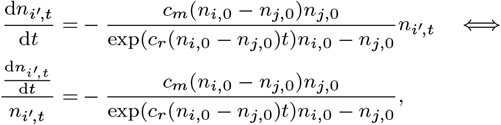

and can thereby be solved by integrating both sides with respect to *t*

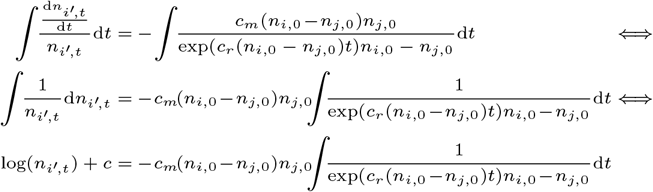

The right hand side integral can be solved by first applying the following substitution *u* = *c_r_*(*n*_*i*, 0_ – *n*_*j*, 0_)*t* where d*u* = *c_r_*(*n*_*i*, 0_ – *n*_*j*, 0_)d*t* ⇔ d*t* = 1/*c_r_*(*n*_*i*, 0_ – *n_j_, 0*)d*u*, resulting in

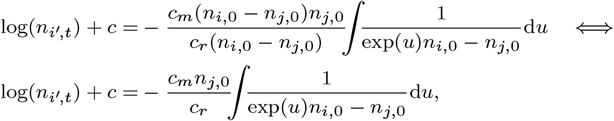

and then apply the following substitution *v* = exp(*u*) where d*v* = exp(*u*)d*u* ⇔ d*u* = (1*/v*)d*v*, resulting in

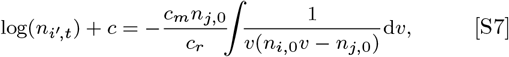

and the right hand side can be simplified using partial fraction decomposition:

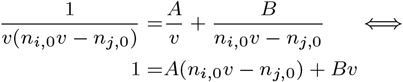

where *v* = *n*_*j*, 0_/*n*_*i*, 0_ gives the solution *B* = *n*_*i*, 0_/*n*_*j*, 0_, and *v* = 0 gives the solution *A* = – 1/*n*_*j*, 0_. Hence,

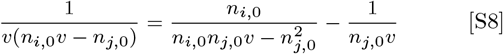

Substituting Eq. S8 into Eq. S7 gives

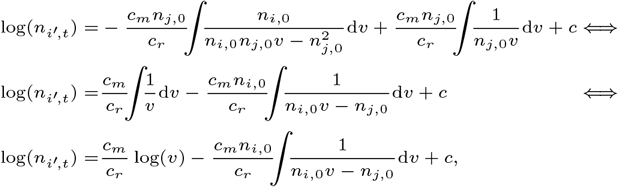

where the integral on the right hand side can be solved with the following substitution *w* = *n*_*i*, 0_*v* – *n*_*j*, 0_ where d*w* = *n*_*i*, 0_d*v* ⇔ d*v* = 1/*n*_*i*, 0_ d*w*, resulting in

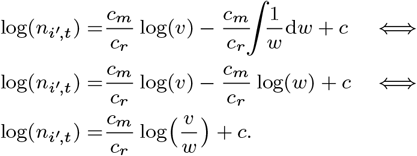

Substitution back *w* = *n*_*i*, 0_*v* – *n*_*j*, 0_, and then *v* = exp(*u*), and then *u* = *c_r_*(*n*_*i*, 0_ – *n*_*j*, 0_)*t* gives

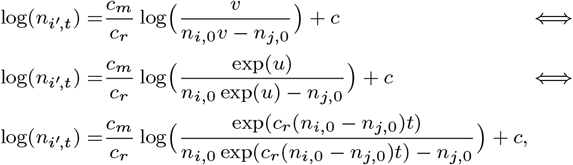

which simplifies to

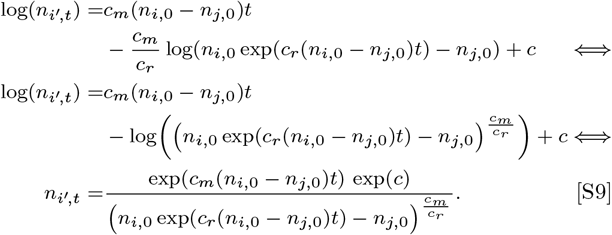

Solving for *c* in Eq. S9 with initial condition *t* = 0 and *n_i′, t_* = *n*_*i*′, 0_ gives

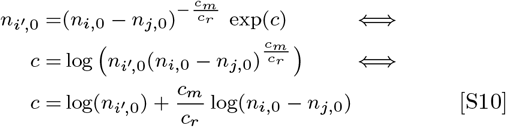

Substituting *c* (Eq. S10) into Eq. S9 gives

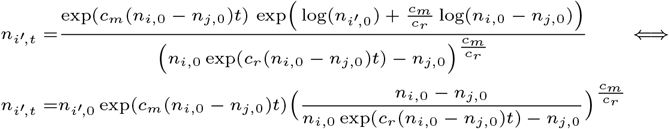

Simplifying this once more gives our explicit expression for the number of gametes of the mutant strategy of mating type *i* at time *t*

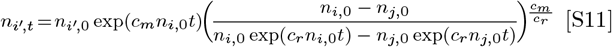

#### 1.4. Solving for gamete fertilisation of a rare mutant strategy under isogamy

For the case of the isogamic constraint, the number of gametes per zygote of mating type *i* with a rare mutant strategy at time *t* decreases according to (Eq. A12)

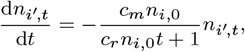

which is a separable differential equation

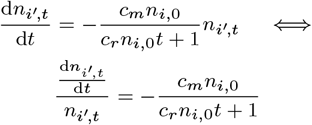

and can thereby be solved by integrating both sides with respect of *t*

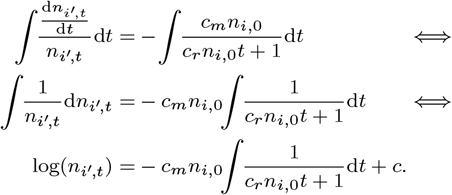

The right hand-side of the integral can be solved with the following substitution *u* = *c_r_n*_*i*, 0_*t* + 1 where d*u* = *c_r_n*_*i*, 0_d*t* ⇔ d*t* = 1*/*(*c_r_n*_*i*, 0_)d*u*, resulting in

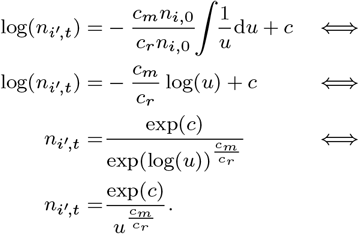

Substituting back *u* = *c_r_n*_*i*, 0_*t* + 1 gives

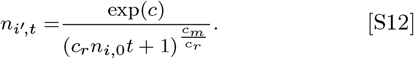

Solving for c in Eq. S12 with initial condition *t* = 0 and *n_i,t_* = *n*_*i*, 0_ gives

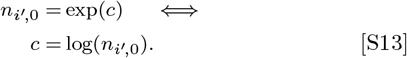

Substituting *c* (Eq. S13) into Eq. S12 gives our explicit expression for the number of gametes of the mutant strategy of mating type *i* at time *t* under isogamy

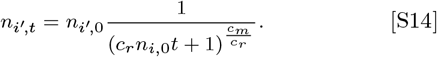

**Fig. S1.**
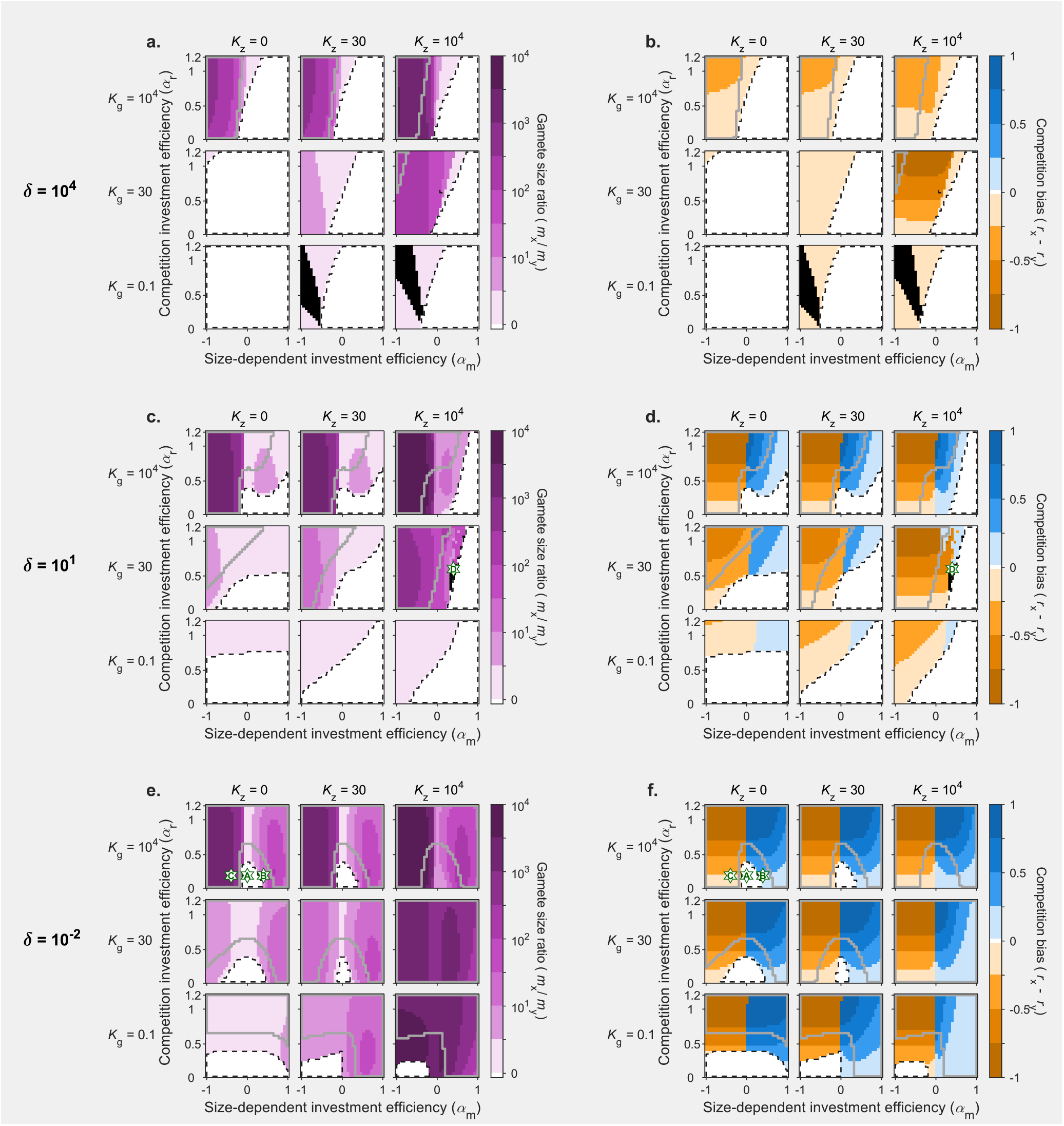
Sex-bias in gamete motility (**b,d,f**) and sexual dimorphism in gamete size (**a,c,e**) at the evolutionary endpoint, as a function of size-dependent investment efficiency *α_m_* (horizontal axis of each subplot) and competition investment efficiency *α_r_* (vertical axis). Size-dependent investment efficiency *α_m_* modulates whether gamete size has a positive or negative effect on the efficiency of energy invested in the competition trait (*α_m_* > 0 or *α_m_* < 0 respectively). The higher the value of competition investment efficiency *α_r_*, the more profitable it is to invest energy into motility. Graphs are shown for three values of gamete density *δ*, which represent variation from a *gamete competition* context (*δ* = 10 000, **a,b**) to a *gamete limitation* context (*δ* = 0.01, **e,f**). Three values of the gamete survival constraint *K_g_* (rows) and zygote survival constraint *K_z_* (columns) are also given. Contour lines correspond to stability analysis of the isogamic singular point. A black dashed line encapsulates the area where isogamy is the expected evolutionary end-point; in the remaining area anisogamy is expected, and a grey contour encapsulates the area where a pseudo-isogamic genetic polymorphism can occur before anisogamy evolves. Coloured shading gives the results from the numerical simulations. Colour intensity expresses the degree of sexual dimorphism in gamete size (**a,c,e**) or investment in competition (**b,d,f**) at the evolutionary endpoint. White represents no dimorphism and deep colours represent strong dimorphism. Stars represent the parameter combinations for which a simulation run is shown in Figure 3, with the upper-case letter (A,B,C,D) referring to the corresponding lower-case letter in the panels of Figure 3.

**Fig. S2.**
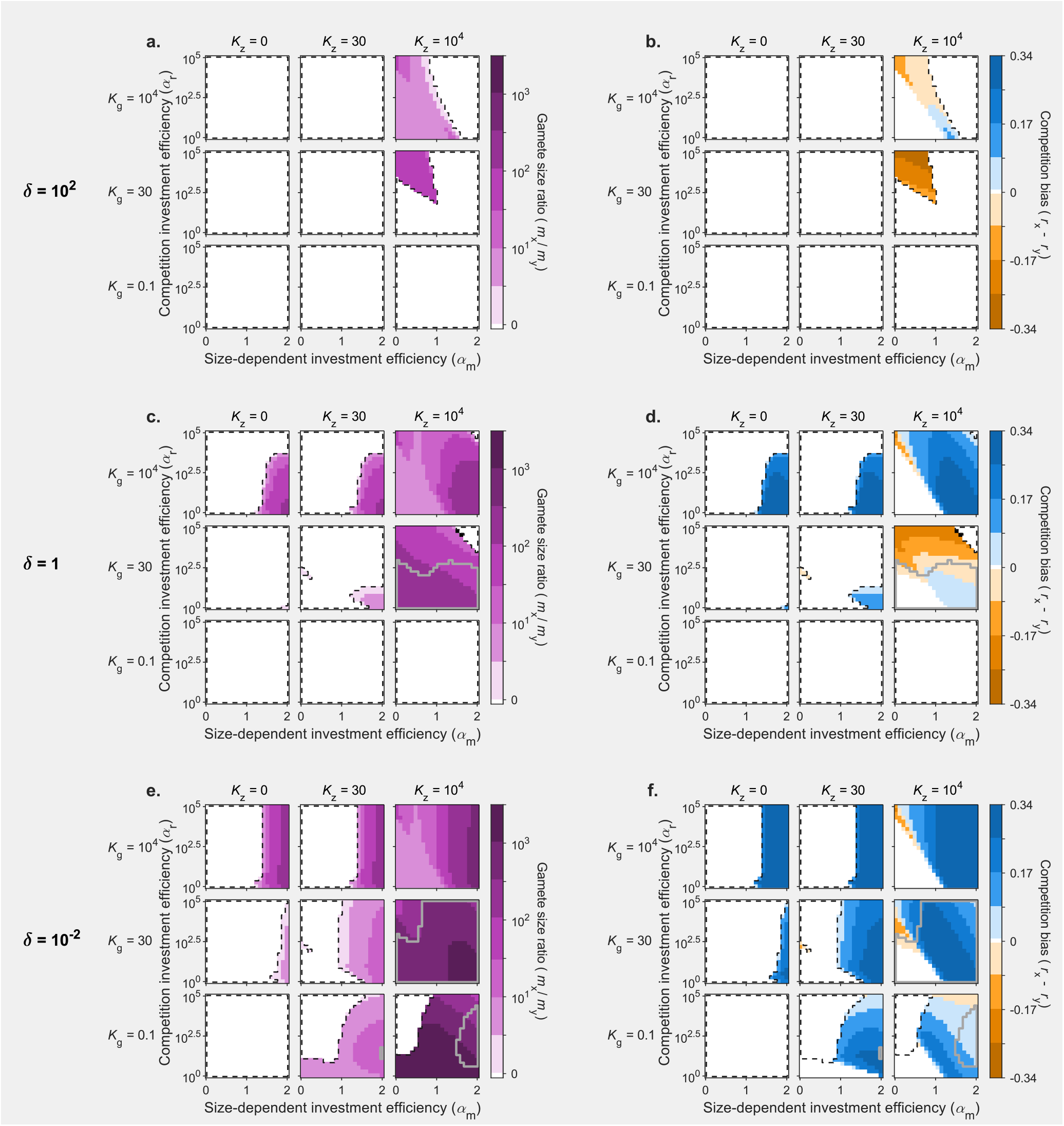
Sex-bias in fusion partner capture (**b,d,f**) and sexual dimorphism in gamete size (**a,c,e**) at the evolutionary endpoint, as a function of size-dependent investment efficiency *α_m_* (horizontal axis of each subplot) and competition investment efficiency *α_r_* (vertical axis). Size-dependent investment efficiency *α_m_* modulates whether gamete size has a positive or negative effect on the efficiency of energy invested in the competition trait (*α_m_* > 0 or *α_m_* < 0 respectively). The higher the value of competition investment efficiency *α_r_*, the more profitable it is to invest energy into motility. Graphs are shown for three values of gamete density *δ*, which represent variation from a *gamete competition* context (*δ* = 100, **a,b**) to a *gamete limitation* context (*δ* = 0.01, **e,f**). Three values of the gamete survival constraint *K_g_* (rows) and zygote survival constraint *K_z_* (columns) are also given. Contour lines correspond to stability analysis of the isogamic singular points. A black dashed line encapsulates the area where isogamy is the expected evolutionary end-point; in the remaining area anisogamy is expected, and a grey contour encapsulates the area where a pseudo-isogamic genetic polymorphism can occur before anisogamy evolves. Coloured shading gives the results from the numerical simulations. Colour intensity expresses the degree of sexual dimorphism in gamete size (**a,c,e**) or investment in competition (**b,d,f**) at the evolutionary endpoint. White represents no dimorphism and deep colours represent strong dimorphism.

